# Developmental variability in cotton fiber cell wall properties linked to important agronomic traits

**DOI:** 10.1101/2024.08.19.607249

**Authors:** Michael C. Wilson, Alexander H. Howell, Anika Sood, Youngwoo Lee, Pengcheng Yang, Elena Yu, Heena Rani, Eileen L. Mallery, Sivakumar Swaminathan, Corrinne E. Grover, Jonathan F. Wendel, Olga A. Zabotina, Jun Xie, Chelsea S. Davis, Daniel B. Szymanski

**Affiliations:** School of Materials Engineering, Purdue University, West Lafayette, IN 47907, USA; Center for Plant Biology, Purdue University, West Lafayette, IN 47907, USA; Department of Botany and Plant Pathology, Purdue University, West Lafayette, IN 47907, USA; Department of Biological Sciences, Purdue University, West Lafayette, IN 47907, USA; Department of Statistics, Purdue University, West Lafayette, IN 47907, USA; Roy J. Carver Department of Biochemistry, Biophysics, and Molecular Biology, Iowa State University, Ames, IA 50011, USA; Department of Ecology, Evolution, and Organismal Biology, Iowa State University, Ames, IA 50011, USA; Department of Mechanical Engineering, University of Delaware, Newark, DE 19716, USA; Department of Materials Science and Engineering, University of Delaware, Newark, DE 19716, USA

**Keywords:** cotton, multiscale phenotyping, CESA, cellulose, morphogenesis, finite element modeling

## Abstract

The economic value of cotton is based on its long, thin, strong, and twisted trichoblasts that emerge from the ovule epidermis. The mature dried fiber cell reflects the outcome of a rapid tapering of the nascent trichoblast, weeks of polarized diffuse growth, followed by a transition to persistent secondary cell wall synthesis. Highly conserved and dynamic microtubule and cellulose microfibril-based anisotropic growth control modules are central to all of these phases. In this paper, we developed novel quantitative phenotyping and computational modeling pipelines to analyze fiber growth behaviors at a daily resolution. We uncovered unexpected variability in growth rate, cell wall properties, and cell geometry across a critical window of fiber development. Finite element computational modeling of fiber growth was used to analyze the instability of cell diameter control and predict how spatial gradients of fiber and matrix material properties can interact to dictate the patterns of shape change. As an initial step toward gaining insight into the molecular orchestration of cellulose biosynthesis, expression profiles of a broad set of relevant genes were quantified across the same developmental timeline and correlated with fiber phenotypes. This analysis identified specific candidate genes that may serve as targets for fiber quality improvement.

## INTRODUCTION

The widely grown cotton *G. hirsutum* (*G.h.*) is the world’s most economically important textile crop species and provides the largest source of renewable textiles ($6.3 billion in raw product, FAOSTAT, http://www.fao.org). This global economy is based solely on the growth and morphogenesis of individual fiber cells that emerge from the developing seed coat and generate a spinnable fiber cell dominated by the properties of crystalline cellulose in the cell wall (Martinez-Sanz, et al., 2017). Given recent advances in knowledge and technologies to analyze cell shape control (Bidhendi and Geitmann, 2018; Gu and Rasmussen, 2022; Yanagisawa, et al., 2022), there are numerous opportunities to use systems level data and computational modeling of cell shape to guide the engineering of cotton with improved agronomic value. In this paper, the spatio-temporal control of growth and cellulose patterning was quantitatively analyzed at unprecedented resolutions that provide new insights into the complex growth behaviors, cell wall properties, and gene expression transitions that underlie important fiber traits.

Developing cotton fibers employ evolutionarily conserved biomechanical control mechanisms to dictate the rates and spatial patterns of growth. Cotton fibers expand through diffuse growth (Ryser, 1977; Seagull, 1986, 1990; Tiwari and Wilkins, 1995; Yanagisawa, et al., 2022), and the growing cell continually synthesizes and modulates the material properties of a tough outer cell wall that deforms in response to turgor-driven axial and circumferential forces (Lockhart, 1965; Proseus, et al., 2000; Szymanski and Cosgrove, 2009). Polarized growth occurs because a highly organized transverse array of cortical microtubules patterns the motility of cellulose synthase (CESA) complexes in the plasma membrane (Li, et al., 2012; Paredez, et al., 2006) and the elongating fiber cells have a resulting transverse network of cellulose microfibrils (Graham and Haigler, 2021; Seagull, 1986, 1990; Tiwari and Wilkins, 1995; Yanagisawa, et al., 2022; Yu, et al., 2019). These CESA complexes synthesize aligned extracellular microfibrils comprising bundles of 18 individual cellulose chains (Nixon, et al., 2016l; Purushotham, et al., 2020). Strong lateral associations among the aligned microfibrils generate a highly anisotropic cell wall that resists deformation parallel to the fibers and are permissive for cell expansion in an orthogonal direction (Baskin, 2005; Paredez, et al., 2006; Zhang, et al., 2021). Consequently, microfibril arrangement at cellular scales drives morphogenesis patterns in cotton (Seagull, 1990, 1992; Yanagisawa, et al., 2022).

Fiber diameter, an important agronomic trait, is first established during a distinct morphogenetic phase of cell tapering (Applequist, et al., 2001; Stiff and Haigler, 2016). The mechanism appears to be conserved, as aerial trichoblasts in many species maintain transverse microtubules along the cell flank and a microtubule-depleted zone at the cell apex that can drive tapering (Graham and Haigler, 2021; Yanagisawa, et al., 2015; Yanagisawa, et al., 2022). These microtubules pattern transverse cellulose fibers in the cell flanks and random fiber arrangements in the apex. This spatial arrangement of differing wall properties leads to a progressive reduction in cell diameter at the tip as the trichoblast elongates (Yanagisawa, et al., 2015; Yanagisawa, et al., 2022). The actin cytoskeleton is also critical for fiber development. Actin bundles have a net longitudinal arrangement in developing fiber cells and are a critical roadway for long distance intracellular transport (Yanagisawa, et al., 2022; Yu, et al., 2019). Distributed secretion to the cell surface must properly replenish cell wall thickness while the cell expands during growth. Genetic experiments in cotton have identified a large number of cytoskeletal proteins that are required for normal fiber growth and that reinforce the overall importance of the cytoskeleton (Gilbert, et al., 2014; Li, et al., 2005; Qu, et al., 2012; Thyssen, et al., 2017; Zang, et al., 2021).

The rate and duration of cell elongation determines fiber length and partially explains the increased length and higher quality of *G. barbadense* (*G.b.*) fibers compared to *G.h.* fibers (Avci, et al., 2013; Beasley and Ting, 1974; Schubert, et al., 1976). However, it is challenging to determine which cell properties control the rate of cell expansion. Parameters like wall thickness, turgor pressure, and matrix modulus affect growth rate, as identified in computational mechanical models of growing trichoblasts (Yanagisawa, et al., 2015; Yanagisawa, et al., 2022). Cotton fibers appear to modulate turgor during the elongation phase by switching on and off their symplastic connectivity to adjacent seed coat epidermal cells to generate a defined interval of elevated turgor (Ruan, et al., 2001). Alterations in cell wall matrix components such as pectin and hemicelluloses may also influence cell wall stiffness (Avci, et al., 2013; Meinert and Delmer, 1977; Pettolino, et al., 2022); however, the specific polysaccharides that modulate growth rate are not known. At an even more basic level, there are disagreements in the literature regarding the potential role of tip growth in fiber cells (Qin and Zhu, 2011; Yu, et al., 2019), and there is a lack of clarity about the relationship between growth rate and secondary cell wall (SCW) synthesis. It is widely assumed that growth rates slow as the cell transitions to SCW synthesis; however, direct comparisons of growth rate and the timing of secondary cell wall synthesis have not been made. The induction of SCW cellulose synthase subunit genes occurs at ∼16 DPA (Betancur, et al., 2010; Pear, et al., 1996). An early study that analyzed fiber elongation rates points to a decreasing growth rate at ∼11 DPA (Schubert, et al., 1973).

Regardless of whether or not SCW synthesis exerts a primary effect on growth rate, the cell undergoes massive compositional and structural rearrangements following the transition to SCW synthesis (Kumar, et al., 2016; Lyczakowski, et al., 2019; Swaminathan, et al., 2024; Zhong, et al., 2019). SCW CESA gene expression is activated (Betancur, et al., 2010; Haigler, et al., 2009; Pear, et al., 1996), as part of a broader transcriptional rewiring of gene expression in which ∼40% of expressed genes have an altered expression level as the cell transitions to SCW synthesis (Grover, et al., 2024).

This paper is part of a set of analyses that were conducted at daily time intervals— RNAseq (Swaminathan S, 2024), proteomics (Lee, unpublished), and glycomics (Swaminathan S, 2024)—in order to obtain a high-resolution portrait of the molecular profiles that span the elongation to SCW phases of fiber development. The multiscale phenotyping reported in this paper revealed numerous examples of dynamic temporal variability in cell wall thickness, growth rate, and microfibril organization. By integrating microfibril behaviors and shape changes with validated finite element (FE) models of trichoblast elongation, the instability of the system and the biomechanical challenges of maintaining a small cell diameter during a protracted developmental program were revealed. In effect, mechanical modeling highlighted the relationships between cell wall spatial gradients of mechanical anisotropy and fiber apex geometry. Lastly, a cellulose synthesis gene expression module was analyzed as a test case to determine how cotton deploys specific subsets of homologous genes to execute a morphogenetic program that fuels the cotton industry.

## RESULTS

### Growth rate and cell wall thickness analysis

To better understand the cellular control of fiber development, data were aggregated to analyze fiber morphology and cell wall traits from the nanoscale of wall thickness to the micrometer scale of microfibril bundling to the millimeter scale of fiber length. Sampling was designed to test for phenotypic variability at daily intervals across the elongation, transition and SCW developmental regimes of fiber growth (Figure 1A). Boll and fiber measurements taken from triplicate samples followed a similar pattern of progressively increasing dimension with an obvious plateau at ∼15 DPA (Figure 1B and 1C, Supplemental Table 1). The mean maximum fiber length underwent a nearly five-fold change from 5-25 DPA as fibers elongated to ∼40 mm. Fiber length data fit over this extended time interval was not linear, and instead fit well to a logistic growth model (Figure 1C). To estimate the fiber growth rate, the derivative of this fitted logistic growth was calculated at points across the 5-25 DPA interval. The peak at ∼11 DPA has been observed with field grown *G.h.* fibers (Schubert, et al., 1973). The entire cell elongates in a diffuse growth mechanism, meaning that absolute growth rate depends on the current length of the fiber. To account for this, the growth rate was divided by the mean fiber length at each sampling point to estimate a relative growth rate. The fiber cell elongated at relative growth rates of ∼30%/day until 8 DPA, after which the rates decreased steadily to nearly zero by 20 DPA (Figure 1D, Supplemental Table 1). The decreases in growth rate (Figure 1C and 1D) preceded the expected transition to secondary cell wall synthesis at ∼15 DPA. Under our growth chamber conditions, upregulation of several SCW CESA genes (Swaminathan S, 2024) and an associated increases in cellulose content (Swaminathan S, 2024) are detected at ∼17 DPA, respectively (Supplemental Figure 1).

**Figure 1:**
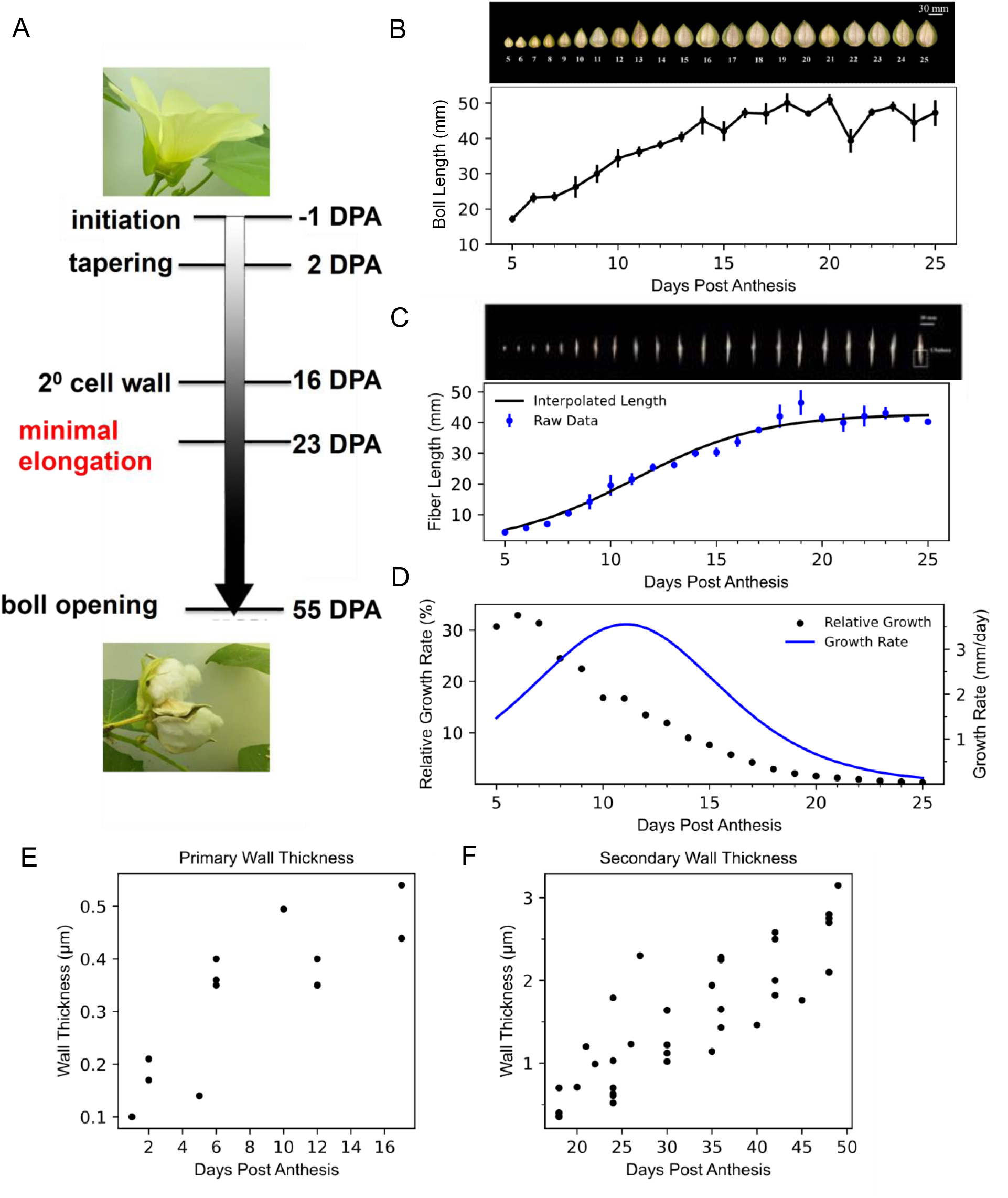
A high temporal resolution analysis of *G. hirsutum* fiber development. (A) Developmental stages of cotton fiber growth, where the white gradient details when fibers are elongating. (B) Daily sampling of cotton bolls and boll lengths (n=3) from 5-25 DPA. (C) Daily sampling of ovules (n=9 for each day) used for measuring the mean maximum fiber length of combed fibers (box at 24 DPA). Mean and standard deviation of the fibers is shown below fitted with a logistic regression curve. (D) The growth rate of fibers was determined from the first derivative of the logistic regression (blue line), and then dividing the rate by mean length of the fibers was performed to measure the relative growth the at daily intervals (black dots). (E) Thickness of the cell wall measured during primary growth and (F) during secondary wall thickening.

Cell wall thickness variability can contribute to cell expansion rates (Yanagisawa, et al., 2015). To determine cell wall thickness variability, published transmission electron microscopy data were mined (Supplemental Table 2) and reliable values were aggregated (Figure 1E and 1F) across developmental time (Avci, et al., 2013; Cao, et al., 2020; Hawkins, 1930; Kljun, et al., 2014; Petkar, et al., 1986; Singh, et al., 2009; Vaughn and Turley, 1999; Wang, et al., 2009; Yanagisawa, et al., 2022). After the tapering phase is completed at ∼2 DPA, wall thickness is maintained at 100-200 nm until 6 DPA, after which the wall thickness appeared to abruptly double and was maintained at ∼400 nm until ∼17 DPA (Figure 1E). Increased thickness due to SCW synthesis was apparent after 20 DPA and the rate of thickness increase appears to be constant until maturity (Figure 1F). The daily changes in cellulose, pectin, and hemicellulose were quantified from *G.h.* bolls grown under identical conditions and were statistically analyzed by ANOVA (Supplemental Figure 1). The proportion of pectin in cell wall fractions fell steadily and mirrored the decreasing elongation. At late stages the relative amount of cellulose and pectin in fiber walls increased abruptly at 24 DPA, indicating a sharp transition to an increased rate of SCW synthesis.

### Microfibril bundling and orientation quantification

The alignment of cellulose microfibrils is a key determinant of fiber shape change and twist (Keynia, et al., 2022; Seagull, 1990; Yanagisawa, et al., 2022). Qualitative analyses of microfibril orientation in *G.h.* fiber reported a shift towards the axial direction at ∼11DPA (Jiang, et al., 2022). Here, we sought to develop a robust and quantitative confocal imaging pipeline to analyze microfibril organization and cell shape at subcellular and cellular scales. Calcofluor-stained fibers were imaged at several overlapping domains with a confocal microscope to generate a montaged image of an extended domain of individual fibers (Figure 2A). The fibers contained an obviously striated pattern of microfibril bundles. The cells were digitally straightened (Figure 2B), and the upper half of the cell was analyzed for shape, fiber orientation (Figure 2C and 2D), and bundling (Figure 2E and 2H). Bundle spacings were consistently about 0.8 bundles/micron until a transition at 15 DPA after which the density of microfibril bundles was reduced to ∼0.6 bundles/µm (Supplemental Figure 4). The non-parametric Dunn’s test revealed a 6-14 DPA grouping and a 15-24 DPA grouping, with significant differences being found almost exclusively in pair-wise comparisons between the two groups (Supplemental Table 3).

**Figure 2:**
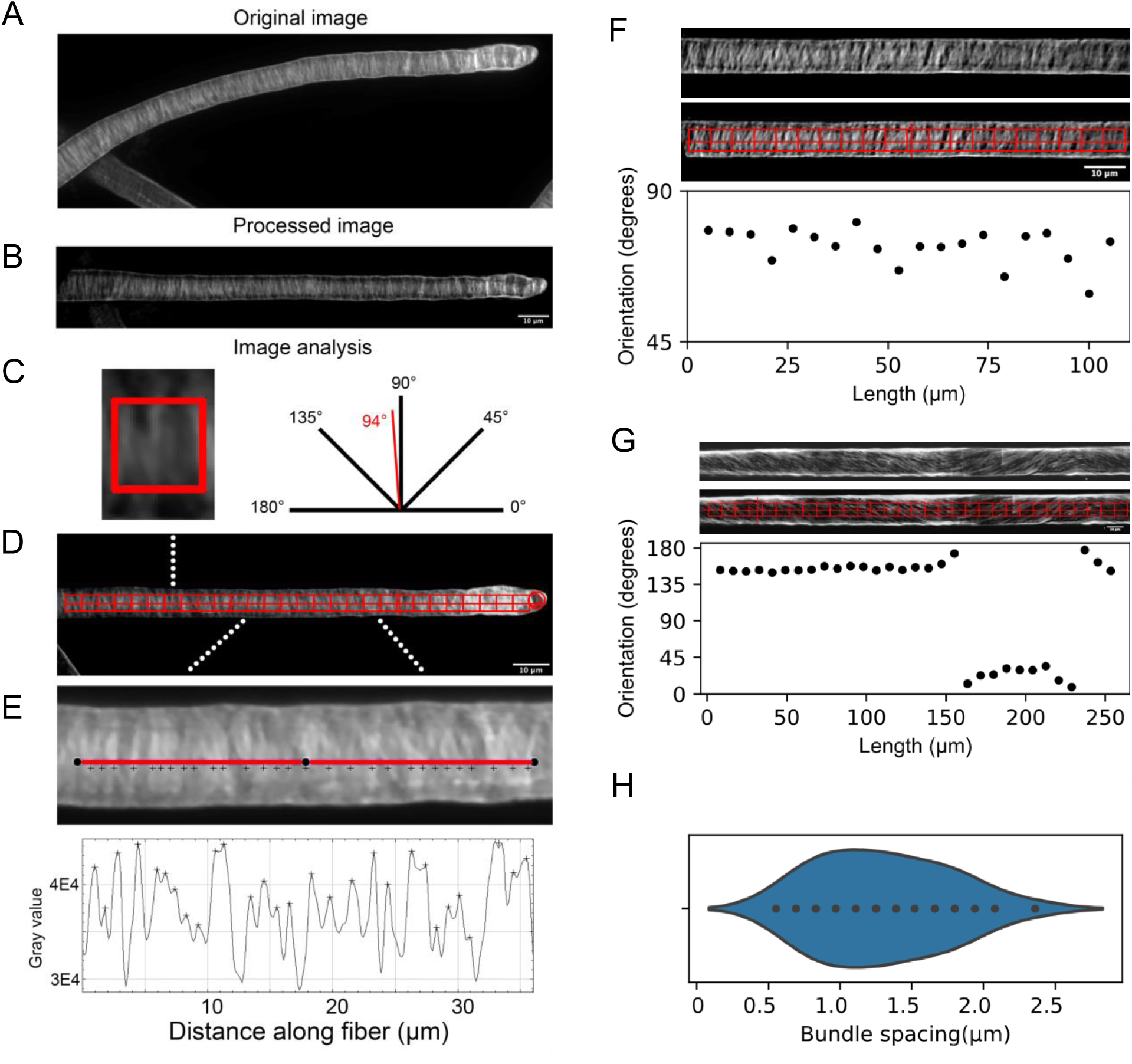
Image processing pipeline for microfibril and cell shape quantification. (A) An example maximum projection of ∼1/2 of the cell volume observed following montaged confocal microscopy of calcofluor-stained cotton fibers. (B) Images were processed to digitally straighten the fiber and subtract background. (C) Schematic diagram defining the angles measured to describe microfibril orientation, with an example from the indicated ROI in (D). (D) The location of the fiber apex, radius of curvature at the tip, and the minimum cell diameter were used to generate an array of ROIs to quantify cell geometry and mean fiber orientation angles. (E) inset of the region outlined in (D) where a line-scan of the intensity profile was used to identify the location of microfibril bundles (individual crosses beneath the line scan). (F) Representative 13 DPA fiber (G) Rare 21 DPA fiber the contains the boundaries in which the handedness of the bundles switches from right to left to right. The measured angles using OrientJ are shown below. (H) Distribution of bundle spacing measurements from the line scan generated in (E).

The image analysis method could also detect transversely oriented bundles along the length of fibers (Figure 2F) and canted helical arrays (Figure 2G) that have been previously reported (Seagull, 1986). The helical arrays were not entirely stable along the length of the fibers and handedness-switching can occur (Seagull, 1986). The cell domains with microfibrils with a fixed handedness were usually large, but in rare instances an entire domain with switched handedness of ∼100 µm could be clearly detected using the OrientJ fiber orientation quantification method (Figure 2G). We consider this to be an estimate of the lower limit of the domain length containing a switched handedness.

The presence of the fiber tip in an image and the biased orientation of the fibers on the slide enabled us to resolve left- and right-handed helical microfibril arrays (Figure 3). The fiber orientation method was applied to quantify the microfibril networks and fiber shapes throughout the 6-24 DPA interval. Images were collected from two different subcellular locations: at random distal fiber locations without a visible apex (Figure 3A, Supplemental Figure 2) and at apical zones that contained an intact fiber tip (Figure 3B, Supplemental Figure 2). Microfibrils had no consistent pattern in the extreme apex but had clearly organized bundles everywhere else. Initial characterizations of microfibril angles entailed calculating the mean orientation angle of all regions of interest (ROIs) within each cell and recording the mean orientation value for all cells in the dataset. The orientation calculation biases towards the dominant features within the ROI and could miss features that are only faintly visible. However, this methodology provides a reasonable estimate. At both subcellular locations, the mean orientations of the microfibrils per cell were dispersed but transversely oriented from 6-14 DPA (Supplemental Table 4). At 15-16 DPA the orientations became more variable in both the apical domain and at distal random locations, and by 20 DPA the orientations shifted to a clearly biased toward a left-handed helical array. The bias was most obvious in the apical domain because the orientation of the fiber was known with certainty; however, all fibers had a biased orientation because of the way they were mounted on the slide (Figure 3C and 3D). The left-handed bias was not stable at later time points. The increased level of variability in mean fiber angle in the cells after 20 DPA reflected mixed populations of either a left- or right-handed dominant arrays in cells, because a clear bimodal distribution of angles are observed when all ROIs are pooled and analyzed in aggregate at each DPA (Figure 3E and 3F). However, the angles of skew relative to the transverse axis were similar in right- and left-handed helical microfibrils (Figure 3G and 3H).

**Figure 3:**
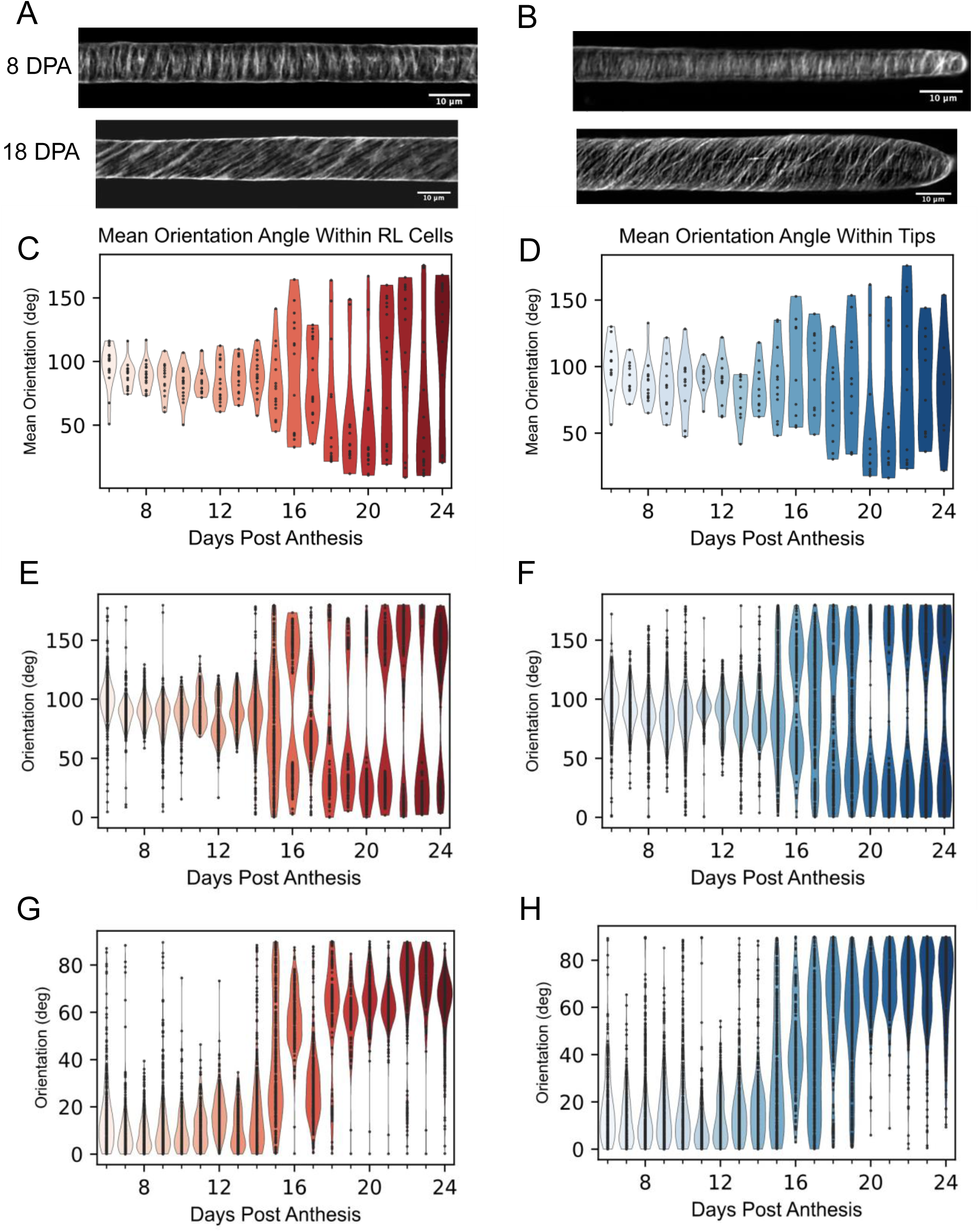
Microfibril organization as function of subcellular location and developmental time. (A) Example images of 8- and 18-days post anthesis (DPA) fibers stained with calcafluor white obtained from random locations along the length of fibers or (B) at the fiber tip. (C) Distribution of mean microfibril bundle angles of cells at random locations from 6 to 24 DPA. (D) Distribution of mean microfibril bundle angles of cells in apical domains from 6 to 24 DPA. (E) Distribution of mean microfibril bundle angles of all individual ROIs measured at random locations from 6 to 24 DPA. (F) Distribution of mean microfibril bundle angles of all individual ROIs measured in apical domains from 6 to 24 DPA. (G) Distributions of mean microfibril bundle angle deviations from transverse measured at random locations from 6 to 24 DPA. (H) Distributions of mean microfibril bundle angle deviations from transverse measured in apical domains from 6 to 24 DPA.

### Tip morphology and fiber diameter control

Several morphological traits were measured from the same confocal images (Figure 4, Supplemental table 5). Fibers of *G.h.* taper to a mean diameter of ∼8 µm (Stiff and Haigler, 2016), but the growth process is noisy with highly skewed distributions of tip geometries (Applequist, et al., 2001; Graham and Haigler, 2021; Stiff and Haigler, 2016; Yanagisawa, et al., 2022). In our dataset, the mean apex diameter usually fell below 10 µm and was variable within each DPA (Figure 4A). Bimodal distributions of apex diameters have been reported for *G.h.* at early developmental stages (Stiff and Haigler, 2016), and certainly, additional sampling would be required to enable reliable statistical analyses of apex curvature across this wide range of DPA. This variability in the minimum cell diameter in the apical domain was reflected in inconsistent morphologies of the fiber apex (Figure 4C, Supplemental Figure 2B, Supplemental Figure 3). “Bulged” fibers with a localized swelling at the apex were frequently observed, as were fibers classified as “re-tapered” based on the assumption that these cells were re-establishing the tapering process from a bulged morphology, but it is possible that this population reflects distal swelling of a highly tapered cell. Aberrant tip morphologies were common across developmental time, as nearly half of the fiber tips deviated from the gradually tapered morphology that is often observed during the tapering process (Graham and Haigler, 2021; Stiff and Haigler, 2016; Yanagisawa, et al., 2022). The imperfect nature of cell diameter control was also reflected in an increased cell diameter at more proximal random locations compared to those in the apical region. The diameter at the proximal end of the fiber was statistically larger than that at the tip for all but 6, 9, and 22 DPA (Figure 4B). The microfibril patterns in swollen tips was highly variable (Supplemental Figure 2) and there was no clear pattern that might predict or explain the local shape defects from the imaging pipeline. Regions of local swelling were examined for differences in bundle spacing, but no extreme local variability was observed (Supplemental Figure 4).

**Figure 4:**
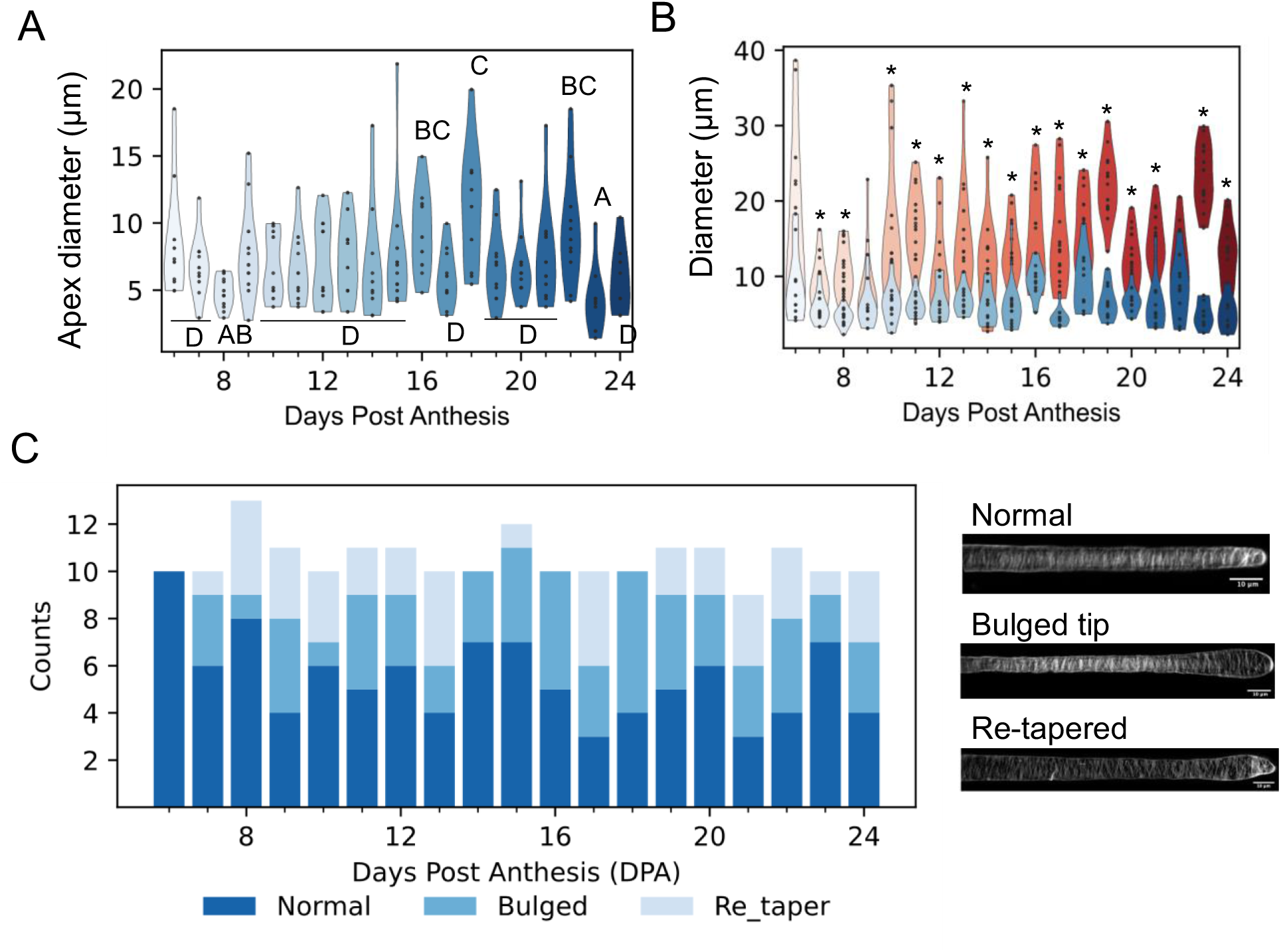
Variability in cell diameter as a function of location and development. (A) Cell diameter variability in the apical domain of growing fiber cells. (B) Comparisons of cell diameter in the apical domain compared with those measured at distal locations (red). Asterisks indicate significant differences between the two subcellular locales. (C) Alternative tip morphologies of the fiber apex as a function of DPA. Tip morphologies were classified as normal, bulged, or re-tapered. Color coded key is shown below panel C.

### Plausible biomechanical causes of apical swelling

FE modeling is an efficient computational tool to predict the types and spatial distributions of cell wall properties that are most critical to generate experimentally observed shape change. FE simplifies geometric or mechanical complexity by representing an object as an assembly of elements with simpler geometries and properties. FE is well suited to problems in plant cell morphogenesis (Bidhendi and Geitmann, 2018) and has specifically been used to analyze a conserved tapering and cell diameter control mechanism in *Arabidopsis* leaf trichomes (Yanagisawa, et al., 2015) and cotton fibers (Yanagisawa, et al., 2022). Here we employed an anisotropic hyperelastic model—the Holzapfel-Gasser-Ogden model (Gasser, et al., 2006; Holzapfel, et al., 2000) similar to Yanagisawa et al., 2015, because it enables the possibility to include variability in the orientation (θ) and dispersion (κ) of cellulose microfibrils that exist in cells and that were measured in Figure 3. Additional key parameters that potently influence cell shape are fiber stiffness (K_1_) which includes fiber density and reflects the extent of material anisotropy, and the matrix modulus (E_m_) (Yanagisawa et al., 2015). In the following simulations we analyzed combinations of variables and their spatial distributions to test for interactions among them in terms of simulated and observed cell shape outputs.

In the first set of simulations, the simulated cells were considered to have uniform material properties over their entire surface (Figure 5). The extreme apical region was defined with isotropic mechanical properties, because it is depleted of organized microtubules (Yanagisawa, et al., 2015; Yanagisawa, et al., 2022) and appeared to lack clearly organized fibers (Supplemental Figure 2). When tested against a range of observed mean fiber angles (θ). The fiber model was seen to axially deform primarily within the flank region but showing more axial deformation near the apex as the fiber bundle orientation deviated from transverse (Figure 5B). However, as reported previously (Yanagisawa, et al., 2022), the fiber globally swelled as cellulose microfibrils deviated from a clear transverse bias (Figure 5C, Supplemental Video 1).

**Figure 5:**
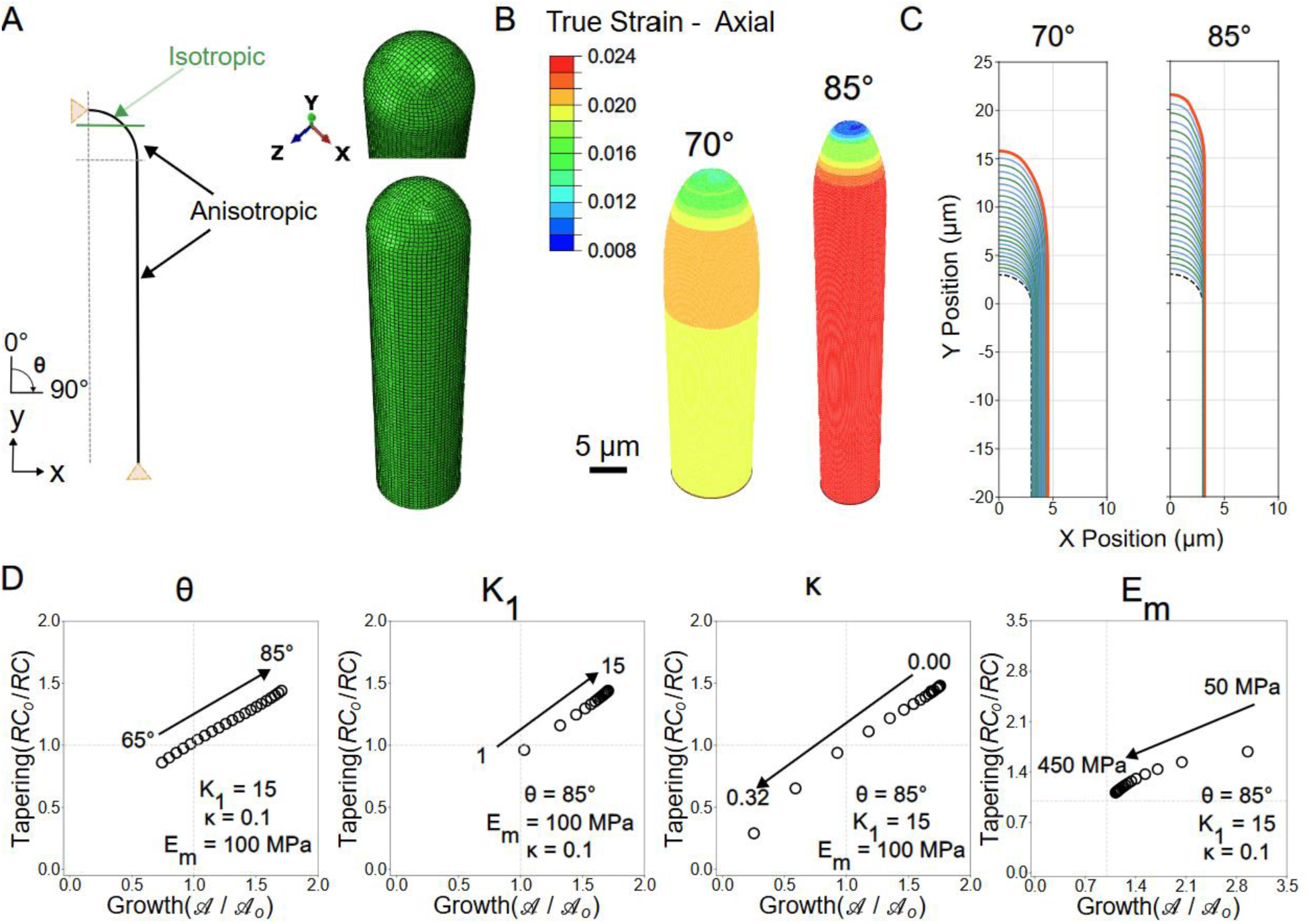
FE simulations and sensitivity analysis of cell diameter and length control for cells with spatially uniform material properties. (A) Design of model geometry and example image of initial simulation mesh. (B) Example logarithmic (true) strain in axial direction on the 25^th^ iteration of simulation with varied fiber angle. K_1_/E_m_ = 15, κ = 0.1, E_m_ = 100 MPa. (C) Example shape of fibers for 25 iterations with varied fiber angle. Black dashed line indicates initial shape. Orange line indicates final shape. Alternating blue and green lines indicate progressive iterations. K_1_/E_m_ = 15, κ = 0.1, E_m_ = 100 MPa. (D) Normalized aspect ratio (𝒜/𝒜_*o*_) and inverse normalized radius of curvature (RC_o_/RC) for fibers with the varied parameters labeled at the top of each panel. (𝒜/𝒜_*o*_) > 1 indicates growth and (RC_o_/RC) > 1 indicates tapering. Open black circle indicates value after 25 iterations.

Both fiber growth and tapering relied on K_1_, κ, E_m_, and θ; as variation in these parameters altered the aspect ratio and global swelling (Figure 5D). Here, we normalize the anisotropic stiffness K_1_ by the isotropic stiffness E_m_ as a relative anisotropy parameter K_1_/E_m._ An increase in θ or K_1_/E_m_ enabled tapering and elongation, but an increase in κ hindered tapering and elongation. Softening of the matrix material enabled growth and tapering when there was a more transverse oriented fiber. The rate of change in growth and tapering was not constant as parameters change, exemplified in the uneven spacing of different model outcomes shown in Figure 5D. As κ approaches 0, θ approaches 90°, K_1_/E_m_ approaches 15, or E_m_ approaches 450 MPa, the amount of geometric variation decreases with constant changes to a parameter. We estimated a value of κ to be 0.024 by aggregating the mean fiber orientation for each ROI from 6-14 DPA. This calculated value presumably underestimated the value, as the distribution within an ROI was not accounted for and each mean angle was biased towards dominant features within an ROI. However, κ existed within an insensitive regime, with minimal changes in growth or tapering up to a value of κ = 0.2 (Figure 5D). For an estimated K_1_/E_m_ of 8 (Yanagisawa, et al., 2015), K_1_/E_m_ also existed within an insensitive regime (Figure 5D). Overall, changes of these parameters could not reproduce the local bulging that was often observed in the apical domain (Supplemental Figure 5).

To explain local bulging, the cell wall was treated as if it had nonuniform properties from the tip to the flank (Figure 6A). Material gradients have been previously examined using finite element analyses in both isotropic elastic and anisotropic viscoelastic cell developmental models (Fayant, et al., 2010; Yanagisawa, et al., 2015). Cotton fibers early in development (<10 DPA) have been shown to contain gradients of pectin epitopes and rupture properties near the tip region (Stiff and Haigler, 2016). We employed two types of mechanical gradients. Because of a higher degree of methyl-esterified homogalacturonan at the apex in hemisphere tips, the region near the apex was treated with a softer matrix modulus (Figure 6B). Second, as cellulose content may be reduced at the fiber apex, the K_1_/E_m_ parameter was also spatially varied. Variation in θ, κ, tip E_m_, and tip K_1_/E_m_ in the presence of a gradient in E_m_ and K_1_/E_m_ (Supplemental Table 6) were still able to produce axial growth and tapering (Figure 6B). Additionally, a gradient in the matrix modulus was able to produce apical swelling; the severity of which was increased when both gradients were used (Figure 6C).

**Figure 6:**
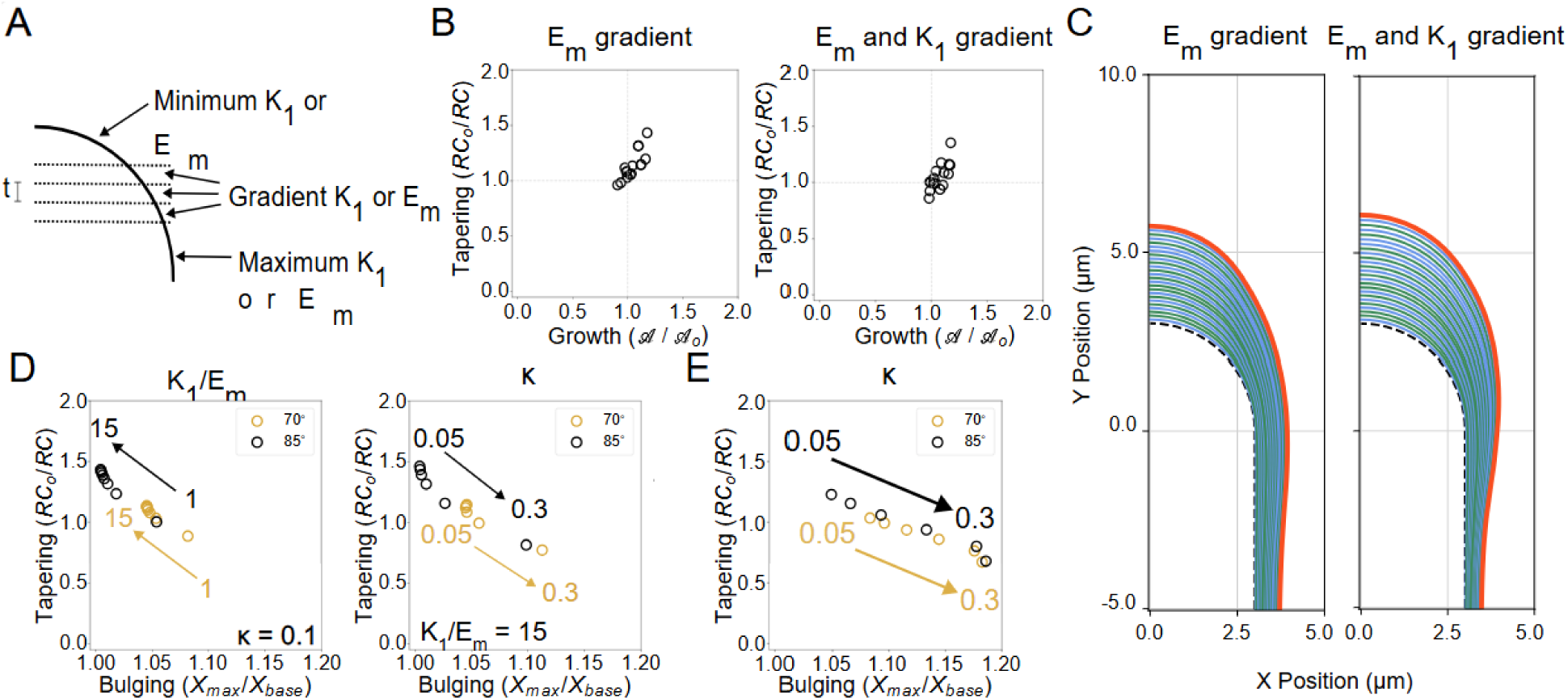
FE simulations and sensitivity analysis of cell diameter and length control for cells with spatial gradients of material properties. (A) Design of gradient with three layers of constant thickness. Initial fiber geometry remained the same as Fig. 5A. (B) Variation in θ, κ, tip E_m_, and tip K_1_/E_m_ with gradient type labeled at the top of each panel. Normalized aspect ratio (𝒜/𝒜_*o*_) and inverse normalized radius of curvature (RC_o_/RC) for fibers with the varied parameters labeled at the top of each panel. (𝒜/𝒜_*o*_) > 1 indicates growth and (RC_o_/RC) > 1 indicates tapering. (C) Example shapes of apical domains for 25 iterations with gradient types listed above each panel. Black dashed line indicates initial shape. Orange line indicates final shape. Alternating blue and green lines indicate progressive iterations. (D) Bulging parameter (X_max_ / X(y = −20)) and tapering parameter (inverse normalized radius of curvature) for fibers with an E_m_ gradient and varied K_1_/E_m_ (left) and κ (right). Open circle indicates value after 25 iterations. (E) Bulging parameter (X_max_ / X(y = −20)) and tapering parameter (inverse normalized radius of curvature) for fibers with E_m_ and K_1_/E_m_ gradients and varied κ. Open circle indicates value after 25 iterations.

Bulging of the shoulder region occurs even in cases of tip tapering, *i.e.* both bulging and tapering values greater than 1 (Figure 6D). However, the degree of bulging was controlled by θ, K_1_/E_m_, and κ. Changes in bulging with κ were seen regardless of whether the gradient was just in E_m_ (Figure 6D) or in both E_m_ and K_1_/E_m_ (Figure 6E). The effect of parameters depends on the value of θ, with different θ influencing both the location and spacing of data in Figures 6D and 6E. Overall, multiple parameters and their spatial distributions can influence cell diameter and length control. Characteristics that resist global swelling also resist local swelling, and because of this, tapering and bulging are inversely related.

### Correlated fiber phenotypes and the expression dynamics of the cellulose biosynthesis network

We next wanted to integrate our fiber growth, wall thickness, and microfibril orientation phenotypes with additional published data on quantitative traits over a similar developmental window (Figure 7). Fiber cells modulate turgor pressure by reversibly gating symplastic continuity with the adjoining seed coat epidermis (Ruan, et al., 2004; Ruan, et al., 2001). A simple interpolation strategy was used based on the five datapoints in Ruan et al., 2001 to generate an estimate of the daily turgor pressure values (Ruan, et al., 2001), assuming that the transitions between measured time points were smooth (Supplemental Table 7). The cellulose, pectin, and hemicellulose levels at the same daily interval have been published (Swaminathan S, 2024). Here, the data were reanalyzed to test for significant differences in cell wall fraction content for both the mass of component per boll and the proportion of each component in the cell wall across developmental time (Supplemental Figure 1). These phenotypes were aggregated with the other informative phenotypes described above and Pearson’s correlation analysis was performed among all possible pairs to group phenotypes that are temporally correlated and potentially related to one another (Figure 7A). Interestingly, the estimated relative growth rate, which is reflective of the diffuse growth mechanism, is most strongly correlated with the rapid initial decrease in the proportion of pectin (Supplemental Figure 1D). Interestingly, the pectin increase at 24 and 25 DPA matches the large increases in cellulose (Supplemental Figure 1A and 1C) and maintains its proportion at these time points, unlike hemicellulose (Supplemental Figure 1D,1F). The observed shift in microfibril angle at 16 DPA was broadly correlated with phenotypes that reflect transition to secondary cell wall synthesis. The microfibril angle correlations were strongest with the amount and proportion of cellulose in the wall (Figure 7A). The estimated turgor pressure profile was not correlated with any other parameter in our dataset.

**Figure 7:**
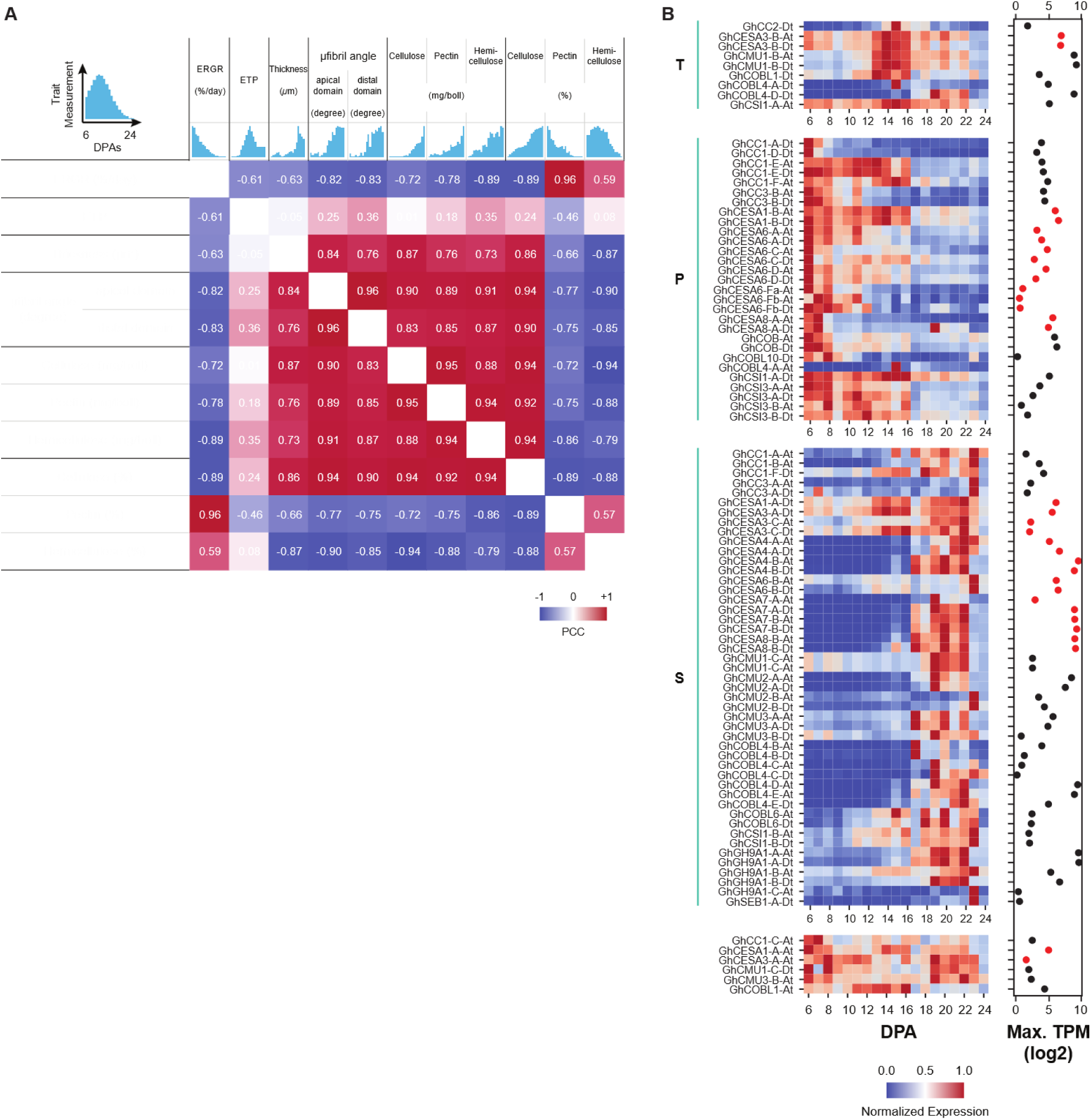
Integrated analysis of fiber phenotypes and candidate *G.h.* genes involved in cellulose synthesis. (A) PCC analysis of fiber phenotypes. (B) Expression variability of a cellulose synthesis gene expression module. Heat map of the expression levels of the fiber-expressed CESAs (highlighted red) and genes known to be involved in cellulose synthesis. Expression level was normalized from 0 to 1 for each gene and organized based on the timing of expression transitions and whether expression was trending down (P), transiently up (T), or trending up (S). P, contains many genes involved in primary cell wall synthesis, T, a transient state that marks a transition to SCW synthesis, S, SCW synthesis. To the right the maximum expression level for the gene across the entire developmental time course is shown, with red dots corresponding to CESA orthologs.

To gain molecular knowledge about which homoeologs might mediate changes in microfibril patterning and cellulose accumulation in developing fibers, we probed a *G.h.* time series RNAseq dataset (Swaminathan S, 2024) for known genes in a cellulose synthesis module that contain CESAs and a variety of other genes encoding enzymes and structural and cytoskeletal proteins (Gu and Rasmussen, 2022; Polko and Kieber, 2019). Cotton orthologs of known *Arabidopsis* genes were identified using inParanoid (Sonnhammer and Ostlund, 2015), and included: orthologs of COBRA (Benfey, et al., 1993; Roudier, et al., 2005; Schindelman, et al., 2001), Glycosyl Hydrolase 9/ KORRIGAN (Nicol, et al., 1998; Urbanowicz, et al., 2007), Cellulose synthase Interacting1 (CSI1), (Gu, et al., 2010) Companion of cellulose synthesis (CC1) (Kesten, et al., 2019), and Cellulose-Microtubule Uncoupling (CMU) (Liu, et al., 2016). The mean expression level in the normalized units of transcripts per million/kB (TPM) are summarized in Figure 7B and Supplemental Table 8. The expression groups, labeled T (transient peak), P (primary) and S (secondary), only partially conformed to an expected sub-functionalization of primary (e.g. CESA1,3,6) and secondary (e.g. CESA4,7,8) cellulose synthases (Tuttle, et al., 2015). For example, numerous CESA1 and CESA6 orthologs have transient peaks of maximal expression within the 6 to 15 DPA window. Along similar lines, and as reported previously for GhCESA4 (Kim, et al., 2011), a subset of predicted SCW CESA4, −7, and −8 orthologs are upregulated to high expression levels after the transition to SCW synthesis (Figure 7B, class S). A proteomic analysis of CESA isoforms across the same developmental window validates that these SCW CESAs are clearly the most highly accumulated enzymes in the CESA family (Swaminathan S, 2024). These expression data reflect a complex utilization of CESA orthologs and instances of homeolog-specific expression.

There were, however, numerous exceptions to this binary division of cellulose synthesis labor. As reported previously (MacMillan, et al., 2017), GhCESA8A homeologs are expressed at their highest level early in development, and proteomic analyses confirm this expression profile at the protein level (Swaminathan S, 2024). Conversely, predicted primary wall synthases GhCESA6B, GhCESA3C, and GhCESA4A homeologs had a maximum expression at 23 DPA. Interestingly, both GhCESA3B homeologs had a transient high level of expression peak centered on 15 DPA (class T). The Dt homeologs of GhCESA3A, GhCESA3C, and GhCESA1A had broad expression patterns with peaks at ∼15 and ∼22 DPA (class S). GhCESA1-A-At and GhCESA3-A-At were part of a loosely organized group (Figure 7B, bottom) that had a more uniform expression across most of the 6 to 24 DPA window.

The expression groups in Figure 7 provides clues about which subsets of additional cellulose biosynthesis genes might work together over time to orchestrate fiber wall assembly. The most striking temporal variabilities were in the GhGH9/KORRIGAN and COBRA and COBRA-like orthologs. The cotton orthologs GhCOB-At/Dt, which are most closely related to *Arabidopsis* COBRA, were the most highly expressed at early stages, with a progressively decreasing expression across DPA. Both COBL4-A homeologs, COBL4-D-At, and COBL1-Dt were strongly and transiently induced during the 13-15 DPA interval. There were 11 COBL4/6/7 genes with a clear induction at or near the transition to SCW synthesis that placed them the “S” expression group. The group was dominated by the 8 COBL4/IRX6 orthologs, including a massive induction from GhCOBL4-E-At from undetectable at 6 DPA to over 400 TPM at 22 DPA (Figure 7B, Supplemental Table 8). All GH9 orthologs had a progressively increasing expression pattern and fell into the “S” group. This does not imply exclusive SCW function because the GhGH9A1-A-At/Dt homeologs were both expressed above 100 TPM early and have extremely high expression levels above 600 by 19 DPA (Figure 7B, Supplemental Table 6). The GH9A1-B-At/Dt pair has a similar profile, but their baseline expression is ∼10 TPM at early stages. A proteomic analysis of protein abundance across the same developmental timeline indicates that these GH9 orthologs mentioned above are the predominant proteins in the fiber cells (Lee, unpublished). The CSI, CC, and CMU genes displayed less temporal variability compared to the GH9 and COBRA orthologs (Figure 7, Supplemental Table 8) and based on the most highly expressed homoeologs we predict the key genes in these orthologous groups to be: GhCSI1-A-At/Dt; both homoeologs for CC1E, F and CC3B, CC1A-Dt, CC1C-At, and the CMU1-B, CMU2-A, and CMU3-A.

## DISCUSSION

Plant cell morphogenesis reflects the output of the complex interactions among turgor-driven stress patterns in the cell wall, cell and tissue geometry, and the material properties of the plant cell wall (Belteton, et al., 2021; Coen and Cosgrove, 2023). Reproducible growth outputs are enabled because the cell somehow can sense geometry and continually tune wall material properties at subcellular scales (Bibeau, et al., 2017; Sambade, et al., 2014; Yanagisawa, et al., 2015). Highly anisotropic diffuse growth and microtubule-based patterning of cellulose fiber properties are used to propagate a self-similar shape in aerial trichoblasts including cotton fibers. This paper included quantitative multi-scale phenotyping at a daily resolution that uncovered unexpected temporal variability in the developmental program. *G.h.* fibers are known to have an unstable cell-diameter control (Stiff and Haigler, 2016; Yanagisawa, et al., 2022). FE modeling and sensitivity analyses of cell shape provide plausible explanations of how diverse combinations of material properties and their spatial distributions influence cell diameter control during fiber development.

### Growth rate reduction precedes secondary cell wall synthesis

Daily sampling of fiber phenotypes and gene expression enabled unprecedented insights into the molecular, cellular, and biomechanical dynamics that underlie fiber development. Boll enlargement and fiber elongation were coupled across a wide developmental window. The profile of fiber elongation was clearly non-linear, and conformed well to a logistic growth function (Figures 1C and 1D). The growth profile mirrored that of fibers analyzed infield-grown plants (Schubert, et al., 1973), with maximal absolute growth rate occurring at ∼10 or 11 DPA. Previous publications strongly support a predominant diffuse growth mechanism for cotton fibers that is accurately quantified using normalized relative growth rates (Graham and Haigler, 2021; Ryser, 1977; Seagull, 1993; Szymanski and Staiger, 2018; Tiwari and Wilkins, 1995; Yanagisawa, et al., 2022) Relative growth rate decreased progressively from 6 DPA forward (Figure 1D). Growth rates of Arabidopsis trichoblasts are dictated in part by intracellular cell wall thickness gradients (Yanagisawa, et al., 2015), and it is generally assumed in that increased cell wall thickness following the transition to secondary cell wall thickness decreases fiber elongation rates. The transcriptional activation of SCW genes occurred from ∼9-14 DPA (Swaminathan S, 2024) and increases in cellulose or cell wall thickness were not apparent until ∼17 DPA (Figure 1E and F, Supplemental Figure 1). Clearly, fiber elongation rates, independent of the method used to calculate them, decrease well before a thicker SCW is detected.

During the tapering and early fiber elongation phases relative growth rates are ∼75-110%/day (Graham and Haigler, 2021). Decreasing elongation rates after 6 DPA coincides with the coalescence of fibers into adherent bundles that adopt a tissue-like growth habit (Singh, et al., 2009). Constrained growth due to mechanical coupling among cells and the bending behaviors necessitated by the crowded environment in the capsule could contribute to a decreased growth rate. Complex interactions among cell wall thickness, matrix stiffness, and turgor pressure modulation affect growth rate in *Arabidopsis* trichoblasts (Yanagisawa, et al., 2015). It is possible that temporal variability in the cotton fiber wall composition (Avci, et al., 2013; Swaminathan S, 2024) and turgor pressure (Ruan, et al., 2001) also influence growth rate. However, the precise molecules that influence the rate of axial expansion are not known.

### Cell diameter instability at the apex and cell flanks

The overall patterns of shape change in the fibers are relatively simple as the cell interconverts between anisotropic growth, radial swelling, and tapering at the apex. Image data (Figure 4, Supplemental Figure 3) indicate that the geometry of the growing fiber is unstable both at the apex and at random locations along the fiber length. (Stiff and Haigler, 2016; Yanagisawa, et al., 2022). Radius of curvature control at the apex appears to be mediated by a persistent apical microtubule-depleted zone that gives rise to an apical cell wall patch in which the cellulose fibers are randomly organized and the wall has isotropic properties (Yanagisawa, et al., 2015; Yanagisawa, et al., 2022). Cellulose imaging of fiber tips failed to reveal any ordered pattern, further supporting the existence of an apical isotropic patch (Figures 2 and 3, Supplemental Figure 2). Ordered pattens of transversely organized microfibrils were evident in growing fibers in the apex flank and at all distal locations (Figure 3). Mean fiber orientation is a critical parameter that influences the degree of radial swelling (Green, 1958; Seagull, 1992). However, we failed to detect obvious differences between normal and bulged domains along the fiber length (Supplemental Figure 4). The FE models enabled us to simulate the contributions of multiple parameters to radial swelling (Figure 5). Variation in fiber orientation (θ), fiber dispersion (κ), and the anisotropic stiffness (K_1_/E_m_) can contribute to swelling behaviors. Therefore, the simplistic measurement of microfibril angle does not capture the full range of properties that can influence radial swelling. In this broader context of material property control, cellular functions like microtubule-dependent cellulose patterning (Paredez, et al., 2006), autocatalytic self-assembly of organized microfibrils (Chan and Coen, 2020), and the distribution and type of secreted matrix molecules (Saffer, et al., 2023) can affect fiber properties and interact to define local growth behaviors. The numerous distinct paths to a resulting radial swelling may explain the widespread occurrence of increased fiber diameter distal to the fiber apex (Figure 4B).

Because of the importance of the apical domain in the establishment and maintenance of fiber diameter we concentrated our FE simulation efforts on this subcellular locale. Fiber morphology at the apex was highly unstable throughout development, with widespread occurrences of apical swelling and apparent attempts to re-taper (Figure 4C). Localized spatial heterogeneity in cell wall composition and material properties has been reported, and in *G.h.* fibers with a large radius of curvature, the apex is a site of preferential rupture following cell wall weakening (Stiff and Haigler, 2016). Therefore, a broad parametric analysis of material properties and their spatial distributions was conducted. In the initial models with uniform mechanical properties over the simulated surface (Figure 5), localized swelling in the apical region did not occur, regardless of the value of mechanical properties (Supplemental Figure 5). Therefore, the geometry of the fiber alone or a change in any individual property is not enough to induce localized apical swelling. Depending on the mechanical property values, spatial gradients in the fiber or matrix stiffness (K_1_/E_m_ and/or E_m_) with softer material at the tip can enable either efficient tapering or induce local tip swelling (Figure 6B). The degree of local swelling depends on the axial-transverse anisotropy. If the softer regions strongly resist radial expansion through increased K_1_/E_m_, increased θ, or decreased κ, fiber elongation can still occur without bulging, reinforcing the importance of the microfibril network in controlling cell geometry. The geometry of the final fiber is also quite sensitive to θ, under uniform (Figure 5D) or gradient (Figure 6D) material properties. This result suggests that patterning a transverse microfibril array through either combinations of microtubule-dependent (Paredez, et al., 2006) and microfibril-dependent assembly (Chan and Coen, 2020) are the most critical activities that enable the cell to propagate a self-similar shape. Our estimates of κ and K_1_/E_m_ from Yangisawa et al. (Yanagisawa, et al., 2015) may indicate relative insensitivity to variation for these parameters. Interestingly, the extent and path of bulging with varied κ changes if the gradient is simply in E_m_ (Figure 6D) or in both E_m_ and K_1_/E_m_ (Figure 6E). The fiber geometry is predicted to be much more senstive to changes in κ if gradients occur in both K_1_/E_m_ and E_m_. The apex may include such material property gradients and may be sensitized to radial swelling.

A key remaining question is how combinations of material properties are tuned to generate consistent patterns of self-similar growth. The strategies of *G.b.* are of particular interest, as its fibers have a consistently lower and less variable diameter, perhaps through managing pectin esterification and cellulose crystallinity in a more effective manner. Accurate measurement of local material properties and a better understanding of how cell wall stress and geometry sensing influences the cortical microtubule array (Li, et al., 2023) are needed to understand the core mechanisms of cell diameter control.

### Developmental transitions and instabilities of cellular scale microfibril bundle networks

At 15 DPA, microfibril ordering becomes more variable within and between cells (Figures 3C-3H). During this transition, the microfibril array transitions toward a bias of left-handed helical arrangement at cellular scales. This timing and pattern correlates with a reported transition to a spiraled pattern of microtubules (Seagull, 1986). Under our growth conditions, the timing of this transition correlates with a massive rewiring of the transcriptome that includes the transcriptional activations of subsets of SCW CESAs (Grover, et al., 2024; Lee, unpublished; Swaminathan, et al., 2024). In *Arabidopsis* leaf trichomes, the cortical microtubule array similarly transitions from transverse to a longitudinal/helical array (Basu, et al., 2005; Zhang, et al., 2005) after the overall shape of the cell is determined at stage 4 and the cell enters a phase of rapid diffuse growth (stage 5) that is rather isotropic in nature (Yanagisawa, et al., 2015). During stage 5, a clear bias toward a left-handed cortical is detected (Keynia, et al., 2022; Sambade, et al., 2014). This left-handed bias in microtubules patterns cellulose fibers in the wall, because upon desiccation the cell twists with the same bias in handedness (Keynia, et al., 2022).

The initial transition to a left-handed bias in bundle orientation may reflect the proposed existence of a specific “winding” layer of cellulose in the developing fiber. However, the left-handed bias is transient, and after 20 DPA, mixed populations of left- and right-handed microfibril bundles occur within and between cells (Figures 3C-3H). It is not known what governs the transitions between transverse, left-, and right-handed microtubule arrays; however genetic evidence in *Arabidopsis* and cotton point to the involvement of tubulin subunits, microtubule-associated proteins, and actin filament nucleators (Buschmann and Lloyd, 2008). Perhaps the initial transition from transverse to helical reflects the attenuation of a cell wall stress-dependent patterning mechanism that could maintain transverse arrays and an apical microtubule-depletion zone(Li, et al., 2023), to some another mechanism in which microtubule structure and cell geometry interact to generate interconvertible spiraled patterns.

In conclusion, this paper reveals previously unrecognized temporal variability and in growth rate, microfibril network organization, and shape control during fiber development. The data provide novel insights into the biomechanical control of fiber elongation, cell diameter, and wall patterning that may determine the degree of twist upon desiccation (Keynia, et al., 2022). These quantitative analyses provide a context to analyze systems of genes and cellular functions that are likely relevant to cotton cell wall biology and trait improvement (Avci, et al., 2013; Pettolino, et al., 2022; Swaminathan S, 2024). Our analyses point to the central importance of cellular-scale patterning of the microtubule and microfibril arrays. Arabidopsis trichomes and cotton fibers share conserved mechanisms of cell diameter control, and wall stress-dependent patterning of the cortical microtubule array is a plausible mechanism to generate an apical microtubule-depleted zone and a global transverse microtubule array (Li, et al., 2023). Further integrating knowledge across trichoblast model systems is likely to accelerate the discovery of key genes and control modules that relevant to specific fiber traits.

## MATERIALS AND METHODS

### Growth conditions

*Gossypium hirsutum* cv. TM1 plants were grown in Classic 2000 pots (Nursery Supplies, Inc) in a growth chamber under a 16/8 day/night cycle at 28 °C in soil (Sungro Professional Growing Mix), perlite (Therm-O-Rock), bark (OrchidSupply.com), and crushed quartzite (Cherry Stone Grit) mixed at a 4:2:2:1 volume ratio. Plants were fertilized at least weekly to saturation with Miracle Gro. Flowers were tagged on the day of anthesis (Day 0) and characterized by their days post anthesis (DPA).

### Cell wall thickness: published data aggregation and measurements

The methodology employed to gather cell wall thickness data encompassed several rigorous steps. Initially, a comprehensive literature review was conducted to source relevant information. Subsequently, Transmission Electron Microscopy (TEM) images were gathered and subjected to meticulous analysis. The measurement of cell wall thickness was executed utilizing ImageJ, a widely recognized image analysis software. To ensure the robustness of the dataset, outliers were identified and eliminated utilizing an 85 percent prediction interval. This statistical approach aided in refining the dataset and enhancing its reliability. Finally, the visualization of the refined data and the generation of conclusive graphs were accomplished through the utilization of R, a statistical programming language. This methodological framework ensured the accuracy and integrity of the collected cell wall thickness data, thereby facilitating comprehensive analysis and interpretation.

### Fiber length measurements and statistical analyses

Whole cotton bolls were excised from the plant on the desired DPA, and each locule was isolated from the boll and placed in 1% v/v Nonidet P-40 (or IGEPAL CA-630) at 80 °C for 15 minutes for locules at 5-14 DPA and 20 minutes for locules at 15-25 DPA. Fibers were rinsed with 10 mM PBS (pH7) and then gently vortexed in 0.025 M HCl for one minute to enable fiber detangling (Stiff and Haigler, 2016). The 0.025 M HCl treatment did not affect the distribution of banded microfibrils along the length of the cells. Seeds were rinsed again with PBS buffer and then stored in PBS and 0.01% thimerosal at 4 °C until use. Mean maximal fiber length measurements were acquired by placing individual seeds on a glass slide, fanning the fibers into a halo around the seed with a stream of buffer, and then manually aligning the fibers parallel to the long axis of the seed. Images of the fibers were collected using an iPhone dual 12MP camera, and the mean maximum fiber length was measured (inset at 24DPA in Figure 1B) using Fiji for three separate seeds in three different locules of three different bolls for a total of nine measurements. To compare cell diameter in the apical domain to those measured in more distal random locations the minimum fiber diameter of each cell was measured. The distributions of values were tested for normalcy using the Shapiro-Wilk test. The Mann-Whitney U test was used in cases in which a dataset was not normally distributed, otherwise the Student’s t-test was used. The number of individuals in each group is defined by the number of high quality images for each DPA and each subcellular location. Tip radius of curvature and local fiber diameter values were obtained from 10 or more cells at all DPA except for 16 and 24 DPA, which had 9 and 8 cells, respectively. Diameter at distal locations was measured from at least 15 cells at all DPA (Supplemental Table 2).

### Cellulose staining and confocal microscopy

To prepare the fibers for confocal imaging, individual seeds were first removed from storage and placed in a petri dish. 10 mM PBS buffer was slowly poured over the seed to fan out the fibers in a halo, and the seed was bisected radially and then longitudinally to reduce fiber density. The chalazal halves were used for staining and imaging. The two halves of the chalaza were placed in 0.5% (w/v) calcofluor white for 10 minutes for staining. Each half was then placed on a glass slide, fibers were fanned out using PBS buffer to create a single plane of fibers, and then they were dissected from the seed and carefully covered with a glass cover slip to minimize crushing the fiber cells. Samples were mounted on an inverted spinning disc confocal microscope (Zeiss Observer.Z1) and imaged using a water immersion lens (63x, 1.2NA, Zeiss) and a Prime 95B sCMOS camera (Photometrics) linked with the Slidebook software (Intelligent Imaging Innovation). Z-stacks of fibers were collected at 0.3 µm intervals using a 405 nm UV laser for excitation. Fiber cells were judged as acceptable for quantification if the cell retained its round shape based on focusing in Z, it did not cross another fiber cell, and the cell lacked artifactual punctate fluorescence or local creases. Fifteen or more images were captured for bolls aged 6-24 DPA at random locations along the length of fibers, and ten or more images of apical fiber domains were collected that had an intact tip (Supplementary Figure 2). A montage function was used for image acquisition when isolated fibers were close to the cover slip in more than one field of view.

### Image processing and analysis

All images were processed for analysis using Fiji (Schindelin, et al., 2012). Because of the fiber alignment protocol, most fibers were aligned with the tips oriented in a single direction. The image stacks were transformed about the horizonal axis to correct for an image inversion at the camera, and the slice keeper tool was used to select the images that captured half of the cell volume closest to the cover slip. Brightness and contrast of the images were adjusted, and image stacks were rotated slightly so the tip was pointing to the right side of the image. The straightening tool was used to artificially generate a straight fiber, and finally the images consisted of projected maximum intensity. Image analysis was performed using a custom-built macro that prompted the user to define the fiber width at the narrowest region of the fiber, the length of the fiber, and a circle at the tip of the fiber to measure the radius of curvature or diameter at the apex. The macro then generated a sequence of square ROIs with measured lengths that were two-thirds that of the minimum diameter and one circular ROI with a radius two-thirds that of the radius of curvature at the fiber tip for images including tips (Supplemental Figure 2C). Using these ROI, the macro generated a .csv file including fiber length, minimum diameter, ROC at tip, and area of the ROI, and for each ROI, skewness, kurtosis, raw integrated density, and normalized integrated density. Finally, using the OrientationJ plugin the orientation of cellulose microfibrils (Figure 2A) and the coherency of the orientation measurements were collected. We failed to detect any developmental variability in the coherency of the microfibril bundle network. Angles were defined with 90 degrees being transverse to the long axis of the cell. To quantify the microfibril angle changes relative to 90 degrees over developmental time, all mean angles of all ROIs were calculated as the absolute value of the deviation from 90 degrees. The allowed the angle deviations of left-handed and right-handed helical arrays that occurred after 15 DPA to be aggregated and quantified over time. To measure bundle spacing, a five-pixel wide line-scan was performed in Fiji along the midline shank. The gray values and their position along the line were exported and then analyzed using a custom Python script that applied the prominence method to distinguish local maxima by applying a 10% prominence threshold. Local maxima were interpreted as a microfibril bundle, and the number of bundles per 10 microns was measured by finding the distance between the initial and final peak and dividing this by the number of bundles along the line-scan.

### Fiber growth rate calculations

We employed a nonlinear least squares method to fit a logistic growth curve to mean maximum fiber length data spanning from 5 to 24 DPA. The logistic growth model given by the following equation was utilized to describe the growth trajectory of the fibers:

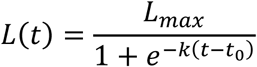

where *L*(*t*) represents the fiber length at time *t*, *L*_*max*_ is the maximum attainable fiber length, *k* is the growth rate parameter, and *t*_0_ is the inflection point indicating the onset of rapid growth. By fitting the observed fiber length data to this model, we obtained the parameters, *L*_*max*_, *k*, *t*_0_. Subsequently, derivatives of the fitted logistic function were computed to obtain the growth rate over the time period. The relative growth rate is further calculated by dividing the growth rate by the fiber length at each DPA (Figure 1D).

### Statistical analyses of cell wall composition from 6 to 24 DPA

Alcohol-insoluble cell wall was extracted from 6 to 25 DPA bolls in triplicates and, further, the pectin and hemicellulose polysaccharide fractions were extracted sequentially as described previously (Avci et al. 2012; Zabotina et al. 2012). In brief, using a sharp razor blade, fibers were separated from the seeds, ground to a fine powder in liquid nitrogen, and the CW was extracted by using organic solvents. The pectin and hemicellulose polysaccharides were extracted sequentially by using 50 mM CDTA:50 mM ammonium oxalate (1:1) buffer (*v*:*v*) and 4 M KOH, respectively. The extracts were dialyzed, dried by lyophilization, and weighed. The final pellet remaining after hemicellulose extraction is defined as the cellulose (mixture of both amorphous and crystalline celluloses), which was also dried and weighed. The three replicates at each DPA were subjected to statistical analysis using R Studio software. The data were initially tested for significance by analysis of variance (ANOVA) and subsequently Fisher’s protected least significant difference (LSD) test value was used to compare treatment means at *p* < 0.05.

### Finite element analysis of cotton fiber development

Finite element analysis (FE) was performed using ABAQUS 2021 (Dassault Systemes SIMULIA). The model used to simulate cotton fiber growth was based on previous FE of single cell systems, namely *Arabidopsis* trichomes—another cell which grows by a diffuse mechanism (Yanagisawa, et al., 2015). Cell growth was modeled through an iterative process using custom python scripts integrated with ABAQUS (Fayant, et al., 2010). In the first step, the cell was drawn as a 2D line structure that was rotated 360° to form a cylindrical shape with a hemispherical cap (Figure 5A). The initial radius of the hemisphere and the cylinder was set to 3 µm, and the height of the cylinder was 20 µm. Then, the cell was pressurized with an internal pressure of 0.2 MPa or 0.7 MPa, based on previous measurements of turgor pressure in developing cotton fibers (Ruan, et al., 2001). Then the coordinates of the deformed cell were extracted from the nodes along the initial drawn line. These coordinates were used to draw the line that was rotated in the subsequent growth step, and the process was repeated for 25 growth steps.

The cotton fiber was modeled with an isotropic region at the tip. While the tip region contains cellulose microfibrils, there is a region of microtubule depletion (Yanagisawa, et al., 2022). As microtubules help in orienting cellulose microfibrils (Paredez, et al., 2006), microtubule absence reduces cellulose microfibril order. Previous experimental work related the size of the microtubule depletion zone (MDZ) with the radius of curvature of the tip region (Yanagisawa, et al., 2022):

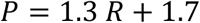

where P is the perimeter of the MDZ (or arc length of the 2D circle) and R is the radius of curvature of the tip. In our simulations, we delineate the tip isotropic zone (TIZ) with a transverse plane located a given distance from the fiber apex (h_TIZ_). This distance can be defined geometrically using the following equation:

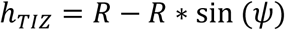

where ψ is defined by the perimeter and radius of curvature.

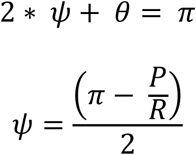

The initial h_TIZ_ is set at 1.21 µm, based on the radius of curvature of 3 µm. In subsequent growth steps, h_TIZ_ is calculated using the radius of curvature from the deformed geometry.

The cotton fiber was treated following the fiber-reinforced Holzapfel-Gasser-Ogden (HGO) model(Gasser, et al., 2006; Holzapfel, et al., 2000), an anisotropic hyperelastic material model. The model considers the isotropic and anisotropic behavior as two additive terms. The HGO model is defined by the matrix modulus (E_m_), a parameter related to fiber stiffness (K_1_), a parameter related to the stiffening of the fibers (K_2_), and the alignment of fiber angles (κ). A κ of 0.33 represents isotropic fiber alignment, and κ of 0 represents fully aligned fibers. To estimate kappa, the distribution of ROI mean fiber orientation from random locations along the cell (Figure 3E) between 6 and 14 DPA was first fit with a von Mises distribution using SciPy (Virtanen, et al., 2020) to determine the concentration parameter (b). Then, the concentration parameter was input into a simplified equation for kappa when calculated as a von Mises distribution (Equation 37 in (Holzapfel and Ogden, 2017)) and solved using Wolfram Alpha. The isotropic region is modeled using κ = 0.33 to represent isotropic cellulose microfibril orientation. In our model, we treated the material as incompressible and assumed a Poisson’s ratio of 0.5. While E_m_ can be linked to physical properties of the pectin matrix, K_1_ does not easily relate to cellulose fiber properties. For the simplest fiber composite models, the contribution of fiber reinforcement to the composite modulus depends on both the fiber stiffness and the fiber fraction. Properties of the cellulose fiber network, like bundling or angle, can be influenced by properties of the pectin matrix (Du, et al., 2020; Saffer, et al., 2023; Yoneda, et al., 2010). Various modes of cellulose fiber deformation can occur as a function of the network organization, altering the observed mechanical behavior (Zhang, et al., 2021). Because of these complexities, it is difficult to directly relate K_1_ to physical behavior of cellulose microfibrils. However, K_1_ can be considered broadly as a kind of anisotropic modulus, with increases in K_1_ increasing the contribution of the anisotropic free energy analogously to how E_m_ increases the contribution of the isotropic free energy. In the original HGO model of arteries, the K_2_ parameter was used to describe fibers which do not initially bear load but bear load upon material extension. In our case, we have kept the value of K_2_ at unity. The mean fiber orientation is defined in ABAQUS. Viscoelastic properties were defined by a 12.5% decrease from initial to infinite modulus and a relaxation time of 6.88 s (Yanagisawa, et al., 2022).

Two boundary conditions were defined. The first prevented the bottommost circular edge of the flank from displacing in the axial direction. The second allowed only axial displacement of the tip apex, preventing fiber bending. The free mesh was defined with a quad-dominated element shape and a seed size of 0.25 µm. Shell elements (S4R) were used. The load pattern was defined by a tabular amplitude with 0% of the applied load at 0 seconds, and 100% of the load applied by 0.005 s. This loading pattern prevented convergence errors in the FE calculation, while still applying the full load over a much shorter time than the relaxation time.

The cell geometry was evaluated through the normalized aspect ratio, and the inverse of the normalized radius of curvature of the isotropic zone. The radius of curvature of the isotropic zone was determined by fitting a circle to the data points with y > (y_max_ - h_TIZ_), where y_max_ is the deformed tip apex position and h_TIZ_ is defined before deformation.

Models were also defined with material gradients from the tip towards the flank. Three transverse planes were drawn at a constant separation from the h_TIZ_ (2 µm). The gradient was defined by three terms: the tip modulus, the flank modulus, and the slice size. The modulus of each section was calculated through a power function:

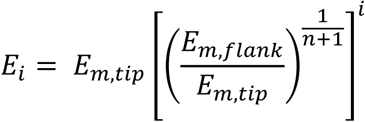

where E_i_ is the modulus for a layer out of *n* layers with the index *i* = 0 corresponding to the tip and progressing towards the flank. E_m,flank_ was set at 400 MPa; E_m,tip_, at 100 MPa or 250 MPa; *n*, at 3. Python scripts are available on GitHUB (GitHUB URL to be provided upon acceptance).

## ACKNOWLEDGEMENTS

The authors would like to thank Dr. Anastasia Desyatova for a base trichoblast FE model and to both Dr. Desyatova and Dr. Thomas Siegmund and for helpful discussions on FE analysis. Artificial Intelligence (ChatGPT and Copilot) was used in the development of code for data analysis and producing graphical elements. This research was supported by the National Science Foundation (NSF) Grant No. 2148122 to C.S.D. and D.B.S. and by Grant No. 1951819 to, J.F.W, O.Z., J.X, and D.B.S.

## SUPPLEMENTAL FIGURE LEGENDS

Supplemental Figure 1. Polysaccharide content of cotton fiber cell walls during development (6 - 25 DPA). Cell wall was extracted from fibers and fractionated into pectin, hemicellulose and cellulose polysaccharides, dried, and weighed. The content of polysaccharides is presented as an absolute content (mg) per boll and also as proportion (%) in relation to other polysaccharides per boll in the graphs.

(A) Absolute cellulose content per boll from 6 to 25 DPA.

(B) Proportion of cellulose content per boll from 6 to 25 DPA.

(C) Absolute pectin content per boll from 6 to 25 DPA.

(D) Proportion of pectin content per boll from 6 to 25 DPA.

(E) Absolute hemicellulose content per boll from 6 to 25 DPA.

(F) Proportion of hemicellulose content per boll from 6 to 25 DPA. All the above data were mean ± SD of three biological replication bolls. Letters (a to i) indicate the significant differences among different DPA bolls (Fisher’s LSD test, *p* < 0.05).

Supplemental Figure 2: All images collected and used for the analyses described in figures 1 & 2.

(A) Images at random locations along the length of the fibers.

(B) Images collected at the fiber tips.

Supplemental Figure 3. Ratio of tip diameter to minimum diameter from 6 to 24 DPA. Ratios between about 0.75 and 1.25 showed the expected, slightly tapered fiber phenotype. Fibers with a ratio around 0.5 generally had a constricted tip region with a swollen section just behind the tip. A ratio greater than 1.5 was indicative of a sock-shaped phenotype whereby the tip was swollen when compared to the more proximal regions of the fiber tip.

Supplemental Figure 3: Ratio of tip diameter to minimum diameter as a function of development. Select images are shown to depict different tip diameters.

Supplemental Figure 4: Local relationships between bundle spacing and cell shape.

(A-D) Region for comparison of bundle spacing within bulged (orange) or normal (blue) regions of fiber in random locations.

(A) (E) Bundle spacing for parts A-D.

(B) (F) Bundle density measured from 6 to 24 DPA.

Supplemental Figure 5: Bulging parameter (X_max_ / X(y = −20)) and tapering parameter (inverse normalized radius of curvature) for fibers with spatially uniform material properties (Figure 5). Open circle indicates value after 25 iterations.

## SUPPLEMENTAL TABLE LEGENDS

Supplemental Table 1: Macroscopic measurements of boll and fiber development in *G.h*.

(A) Boll length from 5 to 25 DPA

(B) Mean maximum fiber length from 5 to 25 DPA

(C) Calculated growth rate and relative growth rate from 5 to 25 DPA

Supplemental Table 2: Reliable cell wall thickness electron microscopy data aggregated from literature.

(A) Cell wall thickness measurements from 1 DPA to 49 DPA.

(B) Cell wall thickness measurements with outliers removed.

(C) Primary cell wall thickness (from 1 to 17 DPA)

(D) Secondary cell wall thickness (from 17 to 49 DPA) with outliers highlighted.

(E) Key for highlighting in table S2A.

(F) Cell wall thickness for *G.b.* from 6 to 49 DPA.

Supplemental Table 3: Dunn’s test for comparisons of bundle spacing with comparisons highlighted among DPAs that are not significantly different.

Supplemental Table 4: Microfibril orientation measurements at tip and random locations from 6 to 24 DPA.

(A) Mean fibril orientation for each cell in images containing a tip. Average values for all cells were calculated as the mean of all ROIs within each cell.

(B) Mean fibril orientation for each cell in images without a tip. Average values for all cells were calculated as the mean of all ROIs within each cell.

(C) Values of fibril orientation for each ROI in images containing a tip.

(D) Values of fibril orientation for each ROI in images without a tip.

Supplemental Table 5: Morphological measurements for cotton fibers from 6 to 24 DPA.

(A) Minimum diameter for each cell in images containing a tip.

(B) Radius of curvature at the fiber apex for each cell.

(C) Minimum diameter for each cell in images without a tip.

Supplemental Table 6: Input (highlighted blue) and calculated (highlighted yellow) values for FE simulations.

(A) Data for Figure 5D with varying θ.

(B) Data for Figure 5D with varying E_m_.

(C) Data for Figure 5D with varying K_1_/E_m_.

(D) Data for Figure 5D with varying κ.

(E) Data for Figure 6B with Em gradient and varying K_1_/E_m_, κ, θ, and tip E_m_.

(F) Data for Figure 6B with Em and K_1_/E_m_ gradient and varying tip K_1_/E_m_, κ, θ, and tip E_m_.

(G) Data for Figure 6D with Em gradient and varying K1/Em and θ.

(H) Data for Figure 6D with Em gradient and varying κ and θ.

(I) Data for Figure 6E with Em and K_1_/E_m_ gradient and varying κ.

Supplemental Table 7: Turgor Pressure interpolation through a first order basis spline connecting experimental data from (Ruan, et al., 2001).

Supplemental Table 8: mRNA abundance profiles of CESA module genes in transcripts per million (TPM) in purified cotton fibers from 6 to 24 DPA.

