## Supplementary figures and images for "Developmental variability in cotton fiber cell wall properties linked to important agronomic traits"

### Supplemental Figure 1

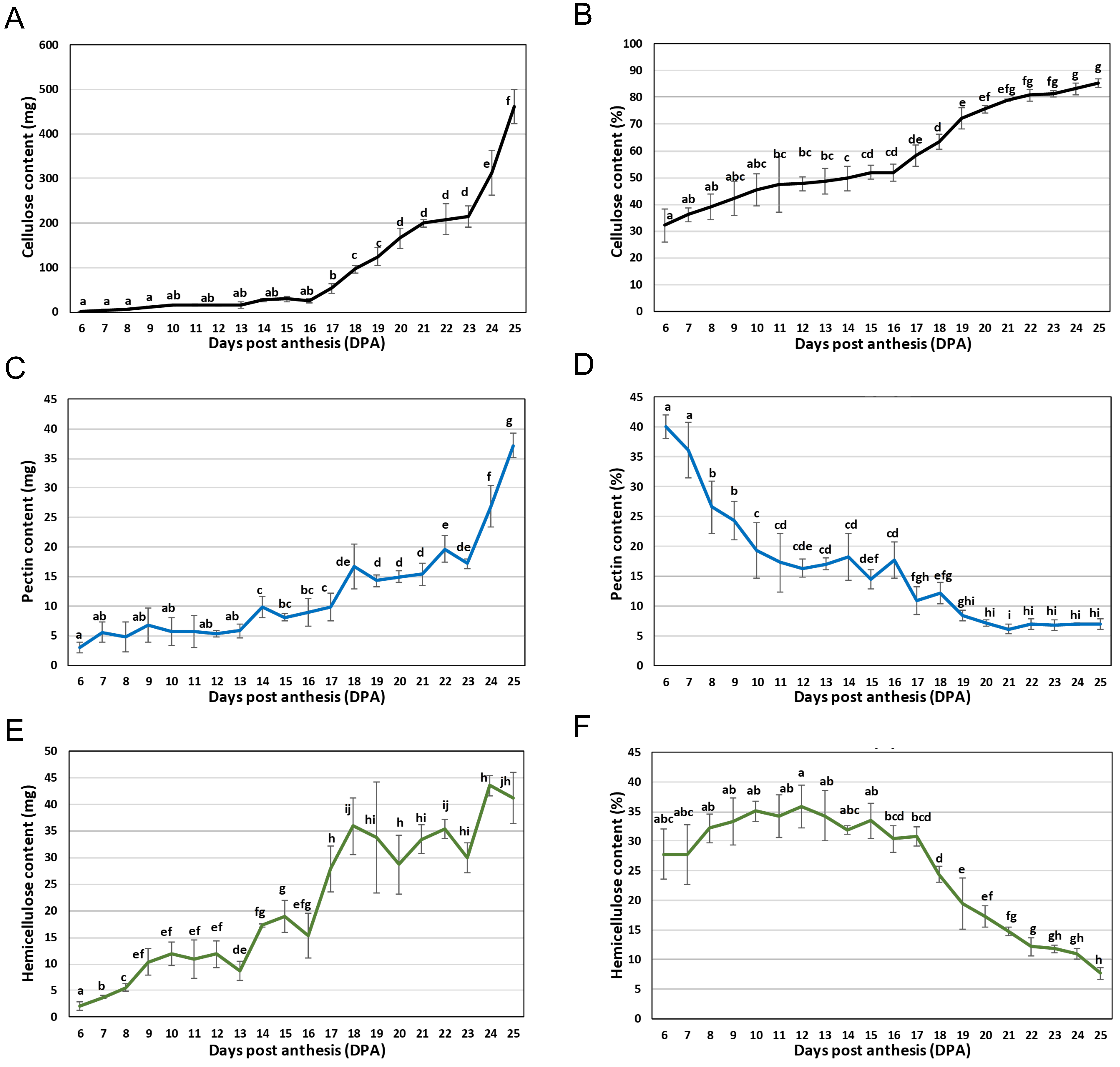

### Supplemental Figure 2

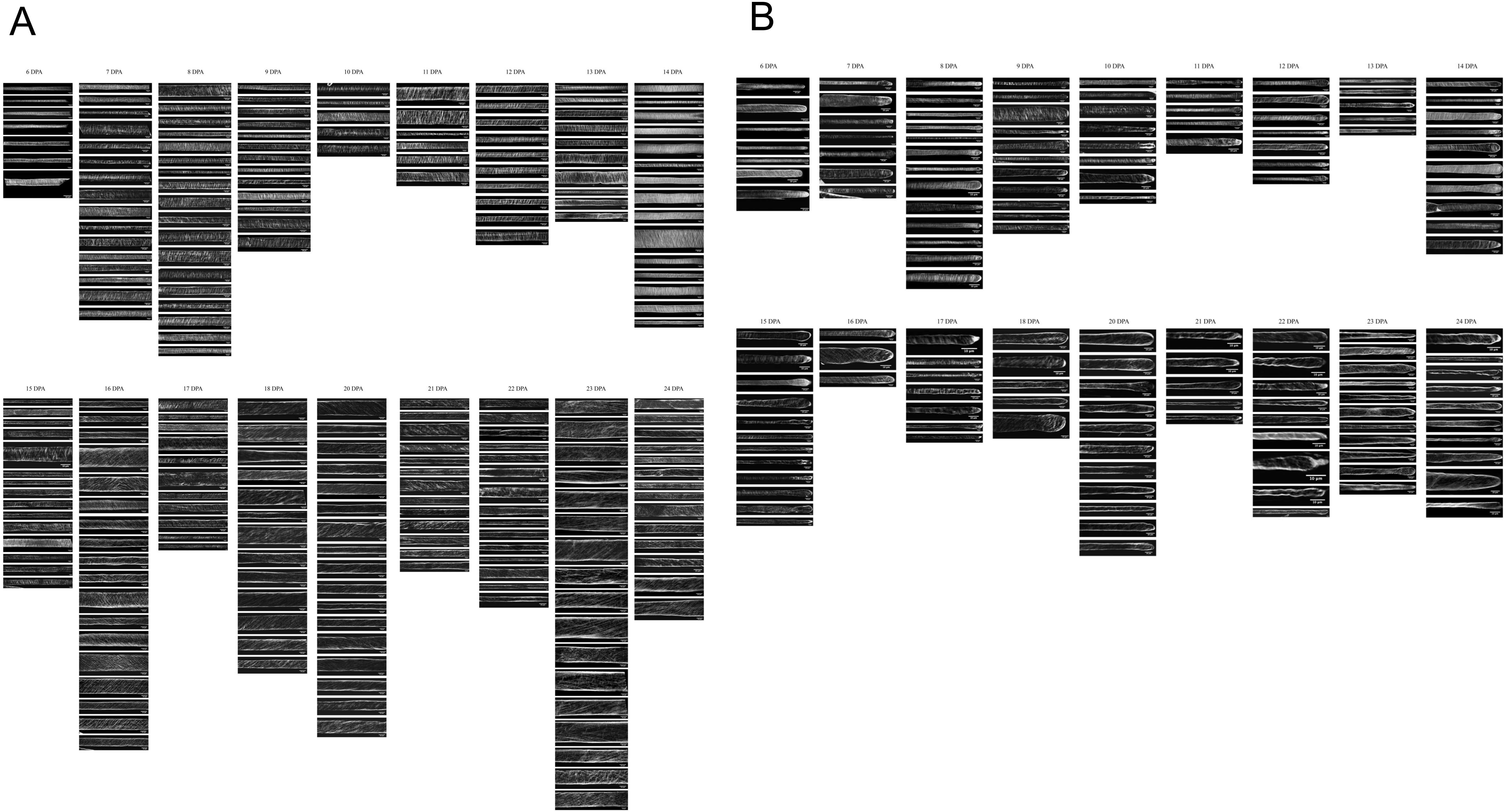

### Supplemental Figure 3

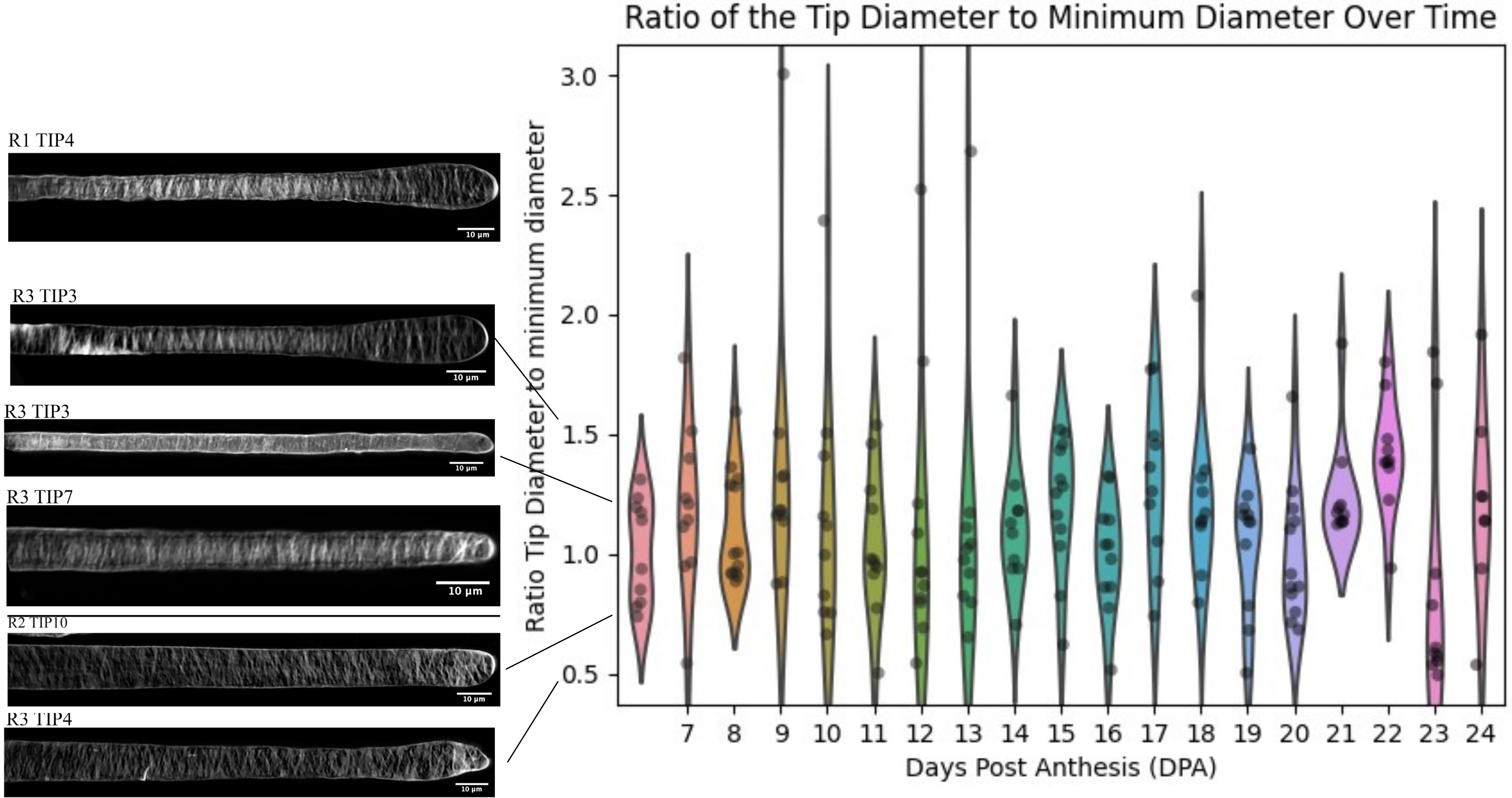

### Supplemental Figure 4

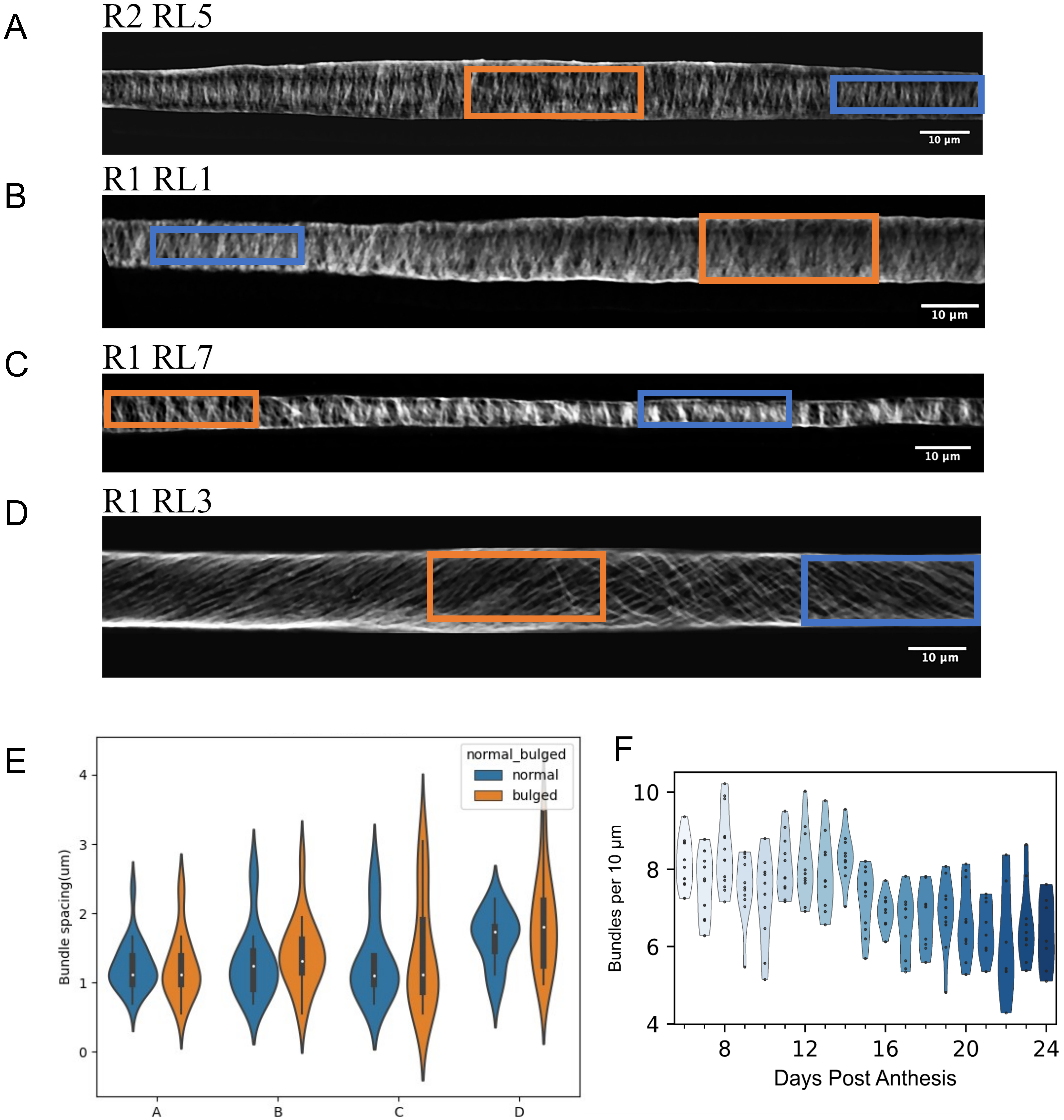

### Supplemental Figure 5

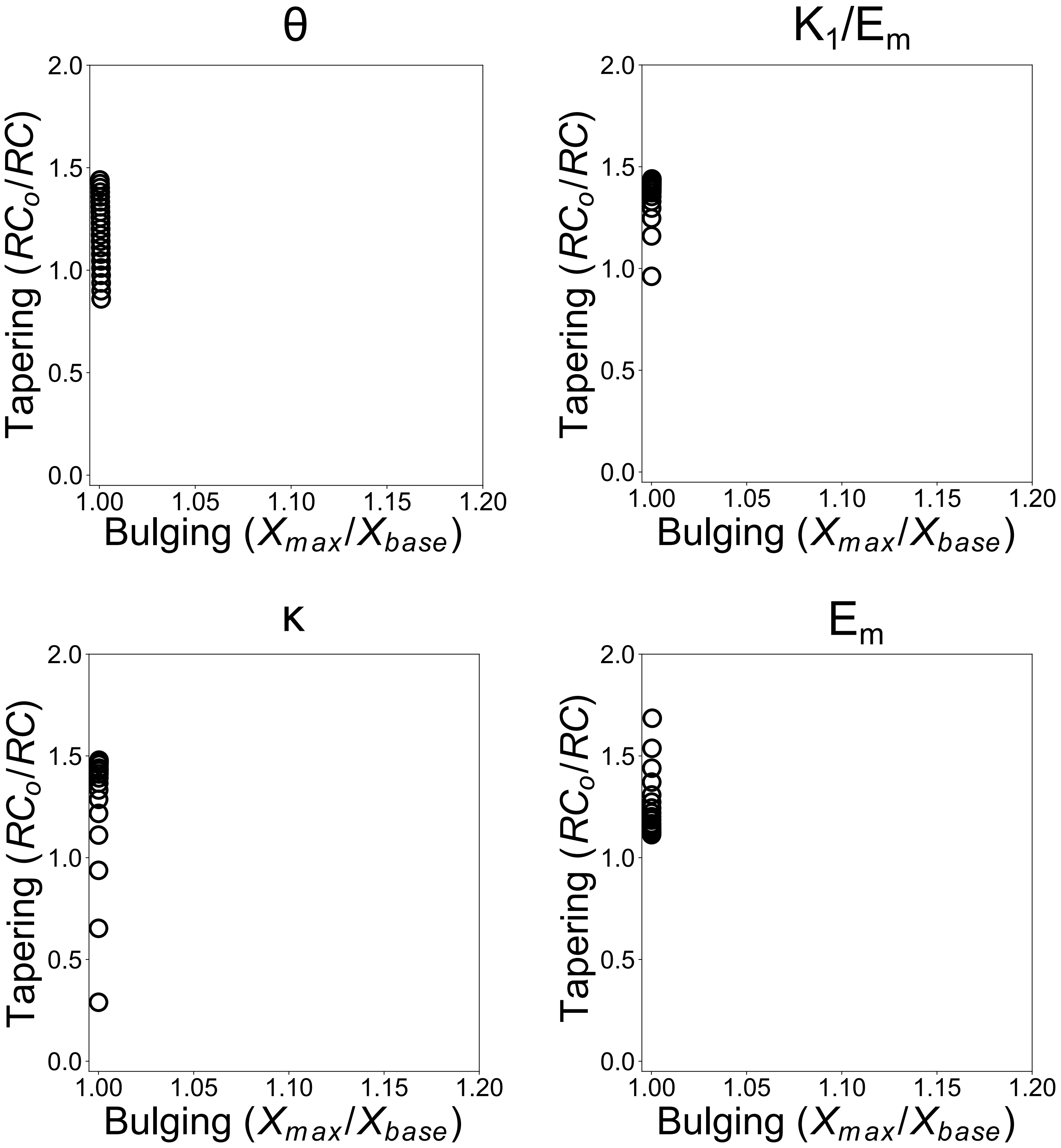

### Supplemental Movie 1

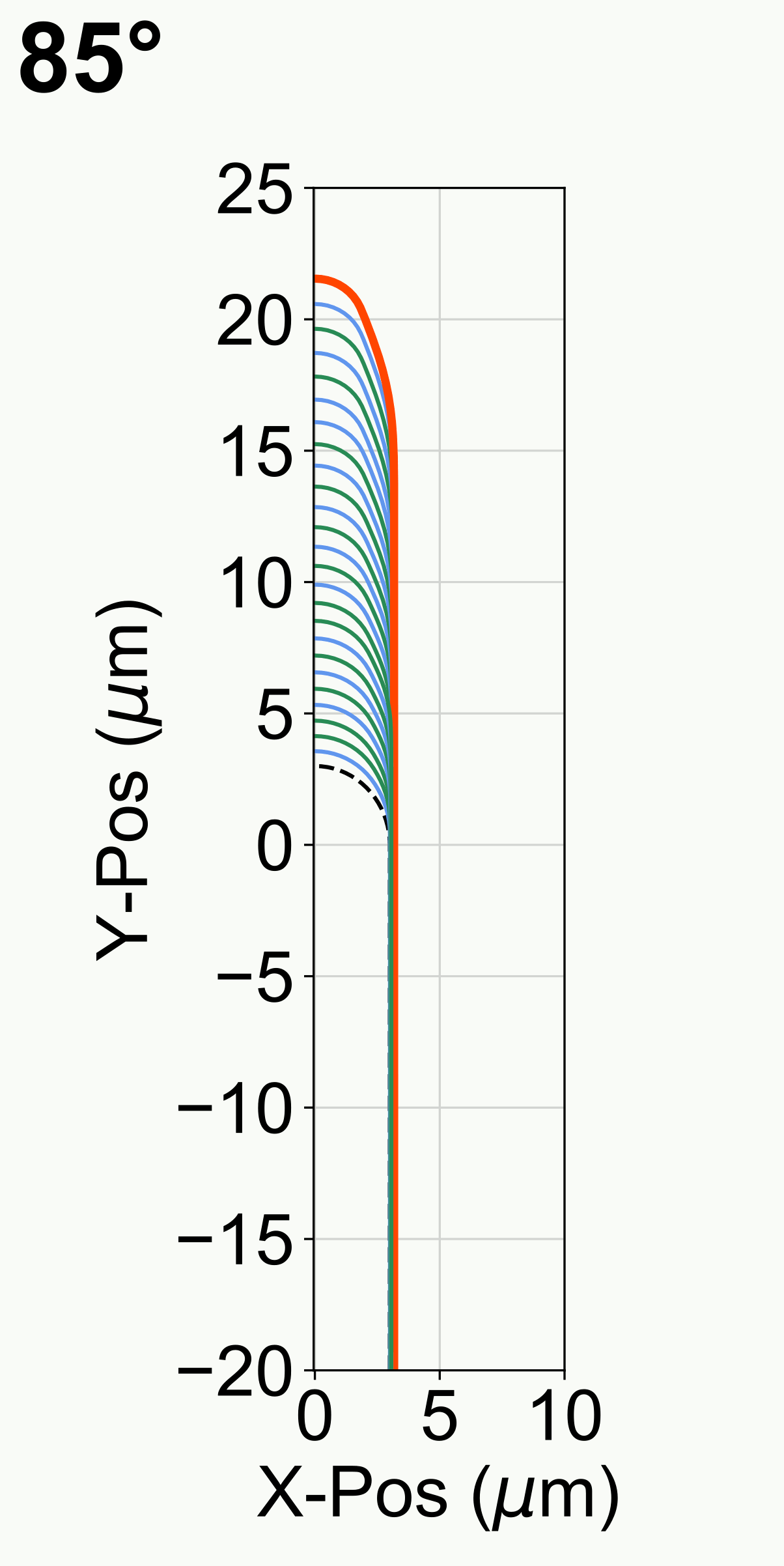
